# T-cell signaling relies on partial CD45-exclusion at sub-micron sized cellular contacts

**DOI:** 10.1101/2025.08.29.673015

**Authors:** Mateusz Kotowski, Daniel F. Heraghty, Markus Körbel, Debasis Banik, Ziwei Zhang, Shekhar Kedia, Bin Fu, James McColl, Bing Li, Joseph Clarke, Martin Fellermeyer, Yuan Lui, Giovanna Bossi, David K. Cole, Haoqi Chen, Ewa Basiarz, Sumana Sharma, Evangelia Petsalaki, Christopher J. Tape, Ben de Wet, Steven F. Lee, Ana Mafalda Santos, David Klenerman, Simon J. Davis

**Affiliations:** MRC Translational Immune Discovery Unit, Weatherall Institute of Molecular Medicine, John Radcliffe Hospital, University of Oxford, Oxford, UK; Radcliffe Department of Medicine, John Radcliffe Hospital, University of Oxford, Oxford, UK; Yusuf Hamied Department of Chemistry, University of Cambridge, Cambridge, UK; Cell Biology and Biophysics Unit, European Molecular Biology Laboratory (EMBL), Heidelberg, Germany; Immunocore Limited, Abingdon, Oxon, UK; EMBL’s European Bioinformatics Institute (EMBL-EBI), Wellcome Genome Campus, Hinxton, UK; Department of Oncology, University College London Cancer Institute, London, UK

## Abstract

How cell contact initiates T-cell activation is uncertain. The local exclusion of the receptor-type protein tyrosine phosphatase CD45 at cell contacts is believed to trigger immune receptor signaling but this is yet to be observed for T cells interacting with authentic cellular targets. Here, quantitative imaging of T cells interacting with tumor cells presenting either native or clinically relevant bi-specific TCR ligands, revealed that they form multiple sub-micron sized ‘close contacts’ with their targets. The contacts were stabilised by the adhesion protein CD2, but efficient ligand detection required both CD2 and integrin ligation. CD45 was excluded from close contacts at the time of ZAP70 recruitment and signaling, but only partially (30− 40%). A single-cell, mass cytometric analysis showed that this change in kinase/phosphatase activity provoked strong T-cell activation and potent cytotoxicity via very small changes in signaling fluxes. Spatial stochastic simulations suggested that the proximal T-cell signaling network is optimised for efficient antigen discrimination in the setting of partial CD45 exclusion. Our work re-frames early T-cell activation as a process initiated by relatively subtle changes in kinase/phosphatase activity acting on small numbers of signaling effectors at minute cellular contacts.

## INTRODUCTION

Cell-cell contact crucially underpins the immune response^1,2^. The cellular architecture of T cells is characterised by finger-like protrusions of their surface referred to as microvilli, which the cells use to interrogate their targets^1,3^. The early contacts thus formed are likely to be dynamic environments where short-lived receptor/ligand interactions occurring on timescales of seconds drive critical decision-making processes. However, the details of these events are poorly understood^1^ because it has been challenging to image early signaling events at genuine T-cell/target cell contacts.

The positioning of the T-cell and target-cell surfaces just 15 nm apart by CD2/CD58 binding, matching the dimensions of TCR/pMHC complexes^4^, ought to lead to the local, steric exclusion of large surface proteins, such as the receptor-type protein tyrosine phosphatase (RPTP) CD45 (>21 nm)^5^, from sites of contact. Indeed, differences in ‘height’ of just 5 nm drive the spontaneous exclusion of large unbound proteins from model contacts created by smaller adhesive complexes^6^. This has important signaling implications, because the local balance of kinase and phosphatase activities will be shifted in favor of the kinases, potentiating receptor phosphorylation^5,7,8^. CD45 exclusion is readily demonstrable for T cells interacting with both glass supported lipid bilayers (SLB)^9^ and SLBs prepared on soft polydimethylsiloxane supports^10^. A substantial body of data also suggests that CD45 exclusion is a pre-requisite of the TCR triggering mechanism. For example, truncating CD45 or extending TCR ligands blocks signaling^11–15^. Moreover, CD45 exclusion alone has been shown to drive receptor signaling in a reconstituted system^16^ and, in some settings, even in the absence of ligands^5^. Similarly, just slowing the diffusion of the TCR at close contacts suffices to initiate signaling^17^. Altogether, these findings imply that TCRs are, essentially, passive structures that only have to be trapped by ligands in phosphatase-depleted close contacts in order to trigger ligand-dependent signaling. However, much of this data was obtained using reductionist, two-dimensional model systems favoring high-resolution fluorescence-based imaging. Target cells are significantly more complicated than the model surfaces and may produce qualitative changes in triggering mechanisms or in the rates of these processes because they are softer and have thicker and more complex glycocalyces^18^. Whether or not RPTPs are actually excluded from contacts between T cells and authentic target cells has not been examined under imaging conditions comparable to those of the model systems.

Here, we studied the early cellular interactions of CD8^+^ T-cells. Using advanced imaging and a bespoke image-analysis platform, we found that T-cells form multiple sub-micron sized contacts with their targets that are stabilised by complexes of the small adhesion proteins CD2 and CD58, leading rapidly to signaling. Integrin engagement profoundly influenced the efficiency of contact formation and CD45 was only partially excluded from the contacts as they formed. A single-cell mass cytometric analysis showed that receptor triggering leading to strong activation and potent target cell killing produces remarkably small changes in downstream effector phosphorylation. Finally, stochastic simulations of TCR signaling suggested that the proximal signaling network is optimised for discriminatory antigen detection in the setting of partial phosphatase exclusion. Discriminatory signaling in T cells therefore relies on relatively limited changes in kinase/phosphatase activity acting on very small numbers of signaling effectors at tiny cellular contacts.

## RESULTS

### ImmTACs as model TCR ligands

We undertook an analysis of receptor signaling by T cells at authentic contacts with cellular targets, using human primary cytotoxic T-cells. We used very thin (<4 µm), adherent U-2 OS osteosarcoma cells as model target cells to facilitate imaging. Previously, we had studied T-cell interactions with SLBs presenting the extracellular domains of the main glycocalyx elements of antigen-presenting cells, *i*.*e*., CD43 and CD45, which project up to 40 nm from the cell surface^9^. Here, we measured the thickness of the U-2 OS cell glycocalyx using spinning-disk confocal imaging (Fig. S1a-d). According to the full width at half maximum values, the glycocalyx extended 250 nm beyond the membrane (Fig. S1e-g). The glycocalyx of U-2 OS cells is therefore significantly thicker than that on the SLBs.

We opted to use pMHC- and TCR-reactive bispecific proteins in the form of ImmTAC®s since they could, in principle, serve as universal TCR ligands for polyclonal primary CD8^+^ T-cells^15,19,20^. ImmTACs consist of two elements: the extracellular regions of TCR-αβ heterodimers linked, *via* the NH_2_ (N)-terminus of the TCR-β chain, to a single-chain variable fragment (scFv) of the anti-CD3ε antibody, UCHT1 (Fig. 1a). The ImmTAC we used binds with picomolar affinity (K_D_ = 55 pM, t_1/2_ = 7.6 h; Fig S2a, Table S1) to the complex of an HLA-A2 molecule presenting a melanoma antigen (the glycoprotein 100 (gp100)-derived peptide YLEPGPVTA)^21^. The UCHT1 scFv binds to CD3ε with nanomolar affinity (K_D_ = 200 nM; Med; Fig. S2b, Table S2). We generated two additional variants of this ImmTAC that were N-terminally-linked with low (K_D_ = 7.9 µM; Lo) and high (K_D_ = 1.8 nM; Hi) affinity anti-CD3ε scFv, rather than the 200 nM form of the scFv (Fig S2c-f, Table S2)^22^. We refer to the three ImmTACs as N-Lo, N-Med, and N-Hi. We first confirmed that ImmTACs can serve as model TCR ligands, *i*.*e*., that they are functional mimics of pMHC. We generated U-2 OS cells presenting ImmTACs bound to gp100/HLA-A2 complexes expressed as single-chain trimers (SCTs, Fig. S3a,b)^23^. First, we examined the relationship between the ImmTAC affinity for CD3ε and potency, using CD25 and CD69 expression as readouts for activation. N-Med was ∼3- and 16-fold more potent than N-Lo and N-Hi, respectively (Fig. 1b and Fig. S3c-e), consistent with previous observations^22,24–26^. To test the effect of ligand length on TCR signaling, we used a variant of the most potent ImmTAC wherein the anti-CD3ε scFv was linked to the C-rather than N-terminus of the TCR-β chain (*i*.*e*., C-Med). This increased the distance between the CD3ε and gp100/HLA-A2 binding sites by ∼7 nm (Fig. 1a) without affecting the affinities of the two moieties (Fig. S2g,h, Table S1,S2), and reduced the potency of the ImmTAC by ∼30 fold (Fig. 1c and Fig. S3f-h). However, extending the gp100-SCT protein by 14 nm by inserting the extracellular region of murine CD4 immediately prior to the transmembrane region of the SCT (producing gp100-SCT_Ext_; Fig. 1a; Fig. S3i), reduced the potency of N-Med by only ∼10-fold (Fig. 1d,e and Fig. S3j-l). This suggests that the different potencies of N-Med and C-Med are not fully explained by length differences alone. Differences in co-receptor engagement (Fig. 1f) explained the remaining difference in potency between the N-Med and C-Med constructs. Mutating the CD8 binding site of gp100-SCT (SCT_KA_, K_D_ > 10 mM; Fig. S3m) reduced the activity of N-Med ∼4.3 fold but did not affect the activity of C-Med (Fig. 1g,h and Fig. S3n-p)^27,28^. The potencies of N-Lo and N-Hi were also reduced (by 6.8-fold and 1.2-fold, respectively (Fig. 1g,h and Fig. S3n-p). Enhancing the affinity of CD8 for the gp100/HLA-A2 SCT by swapping human for mouse β_2_-microglobulin (SCT_A2Kb_, K_D_ = 10.9 µM; Fig. S3m) only substantially increased the potency of N-Lo, consistent with previous reports for native TCR ligands (Fig. 1g,h and Fig. S3n-p)^27–30^. Collectively, these data suggest that, with respect to their ability to trigger the TCR and to engage the CD8 co-receptor, N-terminal ImmTACs are functional mimics of pMHC, insofar as, like pMHC-based triggering, ImmTAC-based signaling is sensitive to the affinity, dimensions, and co-receptor binding activity of the ligand ^12,25,30^. We selected the N-Med ImmTAC as the model ligand in our imaging experiments owing to its optimal potency.

**Figure 1.**
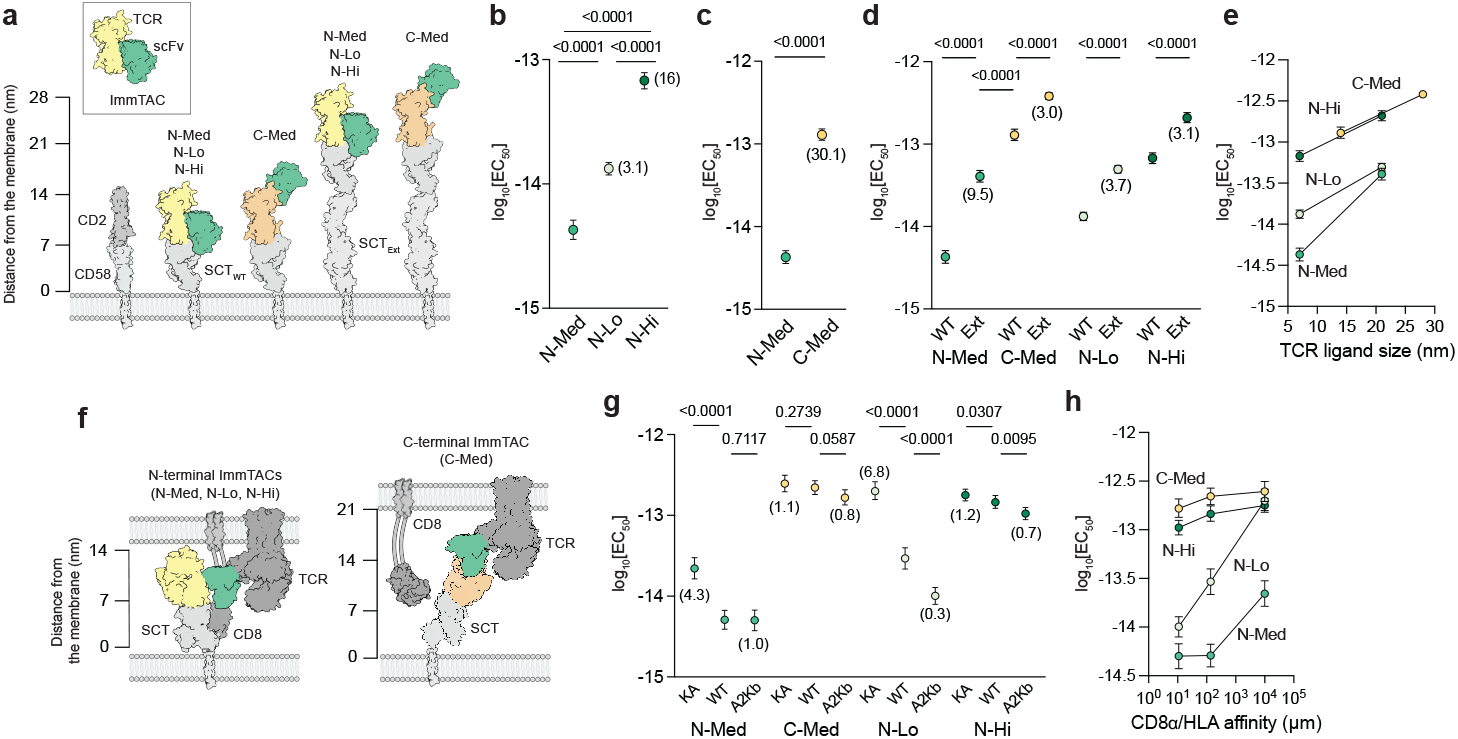
ImmTACs mimic native TCR ligand activity. **a**. Schematic showing N-terminal and C-terminal ImmTACs bound to the wild-type gp100-β_2_M-HLA-A2 single chain trimer (SCT_WT_) and an extended variant of the trimer (SCT_Ext_). SCT_Ext_ was created by inserting the extracellular domain of murine CD4 (measuring ∼14 nm) immediately before the transmembrane region of SCT_WT_. The distance between the antigen and CD3ε-binding sites of the C-terminal ImmTAC (C-Med) is ∼7 nm larger than for the high (Hi)-, medium (Med)- and low (Lo) affinity N-terminal ImmTACs (*i*.*e*., N-Hi, N-Med, N-Lo, respectively). The CD2/CD58 complex is included for comparison. **b**. N-terminal N-Med, N-Lo, and N-Hi ImmTAC EC_50_ values measured in co-cultures of CD8^+^ T-cells and SCT_WT_-expressing U-2 OS cells. Values in brackets indicate the fold shift in EC_50_ values relative to N-Med. **c**. N- and C-Med ImmTAC EC_50_ values measured in co-cultures of CD8^+^ T-cells with SCT_WT_-expressing U-2 OS cells. Values in brackets indicate the fold shift in EC_50_ values relative to N-Med. **d**. ImmTAC EC_50_ values measured in co-cultures of CD8^+^ T-cells with SCT_WT_- and SCT_Ext_-expressing U-2 OS cells. Values in brackets indicate the fold shift in EC_50_ relative to SCT_WT_. **e**. Plot of EC_50_ values versus the approximate axial dimensions of the TCR ligands. **f**. Schematics showing CD8 co-receptor docking to TCR/ImmTAC/gp100-HLA-A2 complexes incorporating either an N-terminal (left) or a C-terminal (right) ImmTACs. The gp100-HLA-A2 SCTs had modified co-receptor-binding sites with increased (SCT_A2Kb_) or decreased (SCT_KA_) affinity for the CD8α subunit relative to SCT_WT_. **g**. ImmTAC EC_50_ values measured in co-cultures of CD8^+^ T-cells with SCT_WT_-, SCT_KA_-, and SCT_A2kB_-expressing U-2 OS cells. Values in brackets indicate the fold shift in EC_50_ relative to SCT_WT_. **h**. Plot of EC50 values versus the affinity of the CD8α subunit for the respective SCT variants. EC_50_ values (b-e) and (g-h) were determined by fitting Hill functions to dose-response curves indicating the fraction of CD69^+^ T-cells (see Fig. S1). Error bars correspond to 95% CI. Co-culture assays were performed in triplicate for three biological repeats. Data was collected using CD8^+^ T-cells from at least two donors. EC_50_ values were compared using an F-test of normalised data. Corresponding CD25 expression data is shown in Fig. S1.

### Scanning-stage microvillar contacts

We first examined the interactions of T cells with their targets in the absence of agonist-induced TCR signaling, in order to capture the pre-signaling behavior of the cells, *i*.*e*., during the “searching” and “scanning” stages of contact formation^9^. Previously, we showed that the accumulation of CD2/CD58 complexes marks the formation of microvilli-mediated close contacts at a model antigen-presenting cell surface^9^. *CD58* gene expression by the U-2 OS cells was eliminated using CRISPR/Cas9, and the cells were transduced with a CD58 construct bearing a C-terminal HaloTag, expressed at the levels of wild-type CD58 (Fig. S4a-e). Hereafter, the cell line expressing wild-type levels of HaloTagged CD58 is referred to as U-2 OS_WT_ cells. The U-2 OS_WT_ cells lack LFA-1 ligands required for T-cell spreading and formation of mature immune synapses^31,32^, and were therefore modified by the introduction of ICAM-1, creating a second cell line that we refer to as U-2 OS_ICAM-1_ (Fig. S4d,e). Polyclonal CD8^+^ T-cells co-cultured with either U-2 OS_WT_ or U-2 OS_ICAM-1_ cells showed no signs of activation measured as upregulation of CD25 or CD69, or as cytotoxicity (Fig. S4f-i). Ensuring that CD58-HaloTag was expressed at physiological levels was critical, however, because its overexpression in U-2 OS_WT+_ and U-2 OS_WT++_ cells led to T-cell activation in the absence of TCR ligands (Fig. S4e,j-l). A ∼26- or ∼340-fold increase in CD58-HaloTag expression (in U-2 OS_WT+_ and U-2 OS_WT++_ cells) increased the fraction of CD69^+^ and CD25^+^ T-cells by ∼35%. Previous experiments with advanced bilayers revealed that supraphysiological CD2/CD58 adhesion produced ligand-independent TCR triggering owing to the formation of large phosphatase-excluded contacts^5,9^.

Primary CD8^+^ T-cell/U-2 OS target cell interactions were imaged using two-color epifluorescence widefield microscopy. The target cells were cultured as contiguous monolayers, restricting T-cell interactions to the apical surface of the U-2 OS cells and avoiding the underlying coverslip-glass substrate. CD58-HaloTag was labelled with Janelia Fluor 549. CD45, which is generally believed to be uniformly distributed over the T-cell surface ^1,33^, was used as a surface marker after labelling it with a fluorescent antibody^1,32^. T cells added to U-2 OS monolayers appeared to ‘glide’ across the U-2 OS_WT_ surface, forming occasional, spatially confined regions of contact marked by transient ‘puncta’ of CD58 accumulation (Fig. 2a; Supplementary Movie 1). These puncta were comparable to the contacts formed by T cells interacting with glycocalyx- or quantum dot-presenting bilayers^1,9^. The number of close contacts detected was smaller than expected, given the high density of microvilli on resting primary CD8_+_ T-cells imaged using electron microscopy (Fig. S4m). In contrast, T cells interacting with U-2 OS_ICAM-1_ cells rapidly formed numerous, well-separated close contacts (Fig. 2b; Supplementary Movie 2). The fraction of T cells forming close contacts within 5 minutes increased four-fold in the presence of ICAM-1 (Fig. 2c). To ensure that they were not artefacts of the chosen focal plane, we used an epi-illumination form of selective-plane illumination microscopy (eSPIM)^34^, which allows visualisation of whole cells, confirming our findings (Figure S4n,o; Supplementary Movies 3,4).

**Figure 2.**
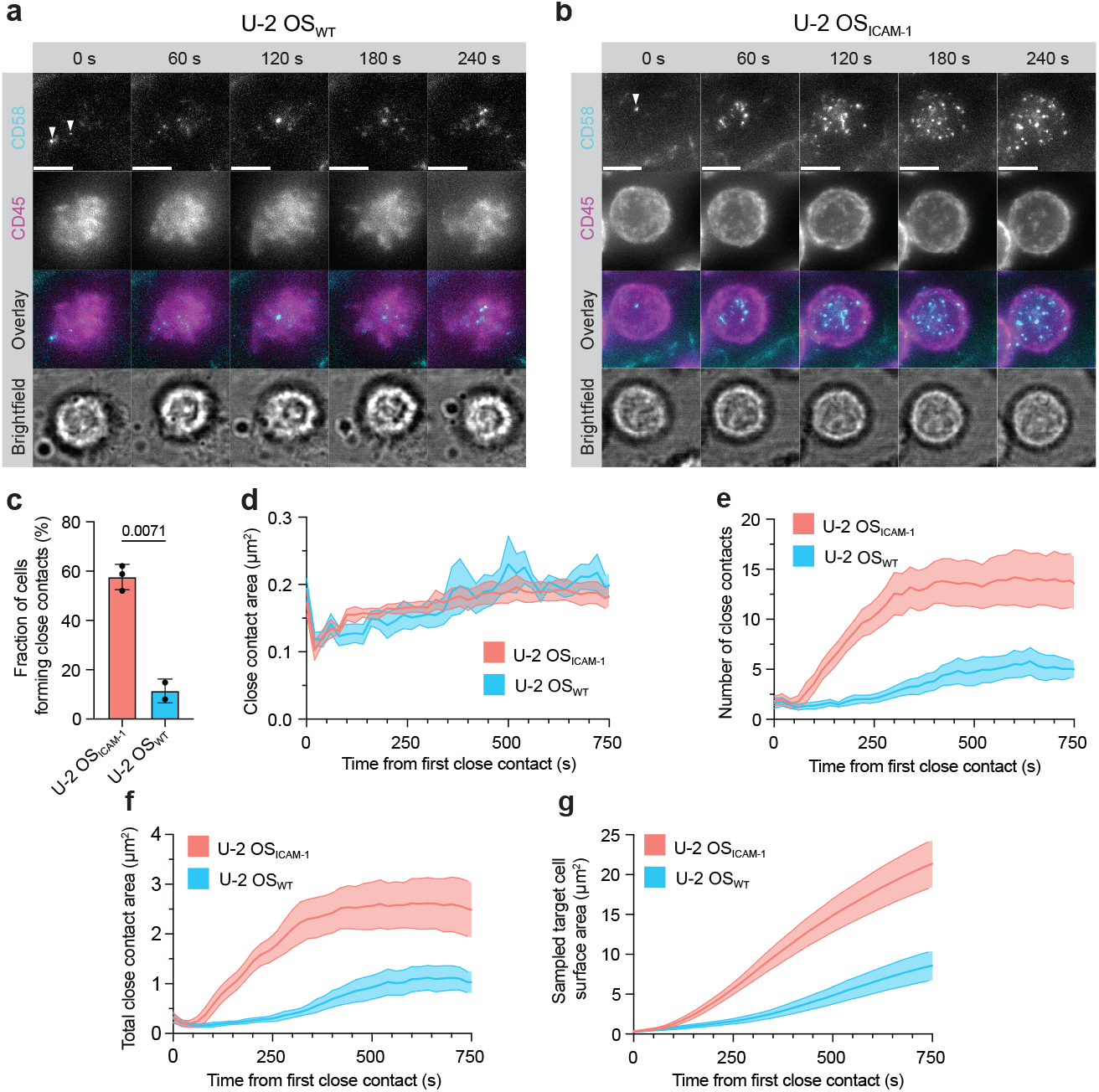
T cells form CD2/CD58-stabilised close contacts with target cells but require integrin engagement to do so efficiently. **a, b**. Epifluorescence-based widefield imaging of CD8^+^ T-cells interacting with monolayers of U-2 OS_WT_ (a) and U-2 OS_ICAM-1_ (b) cells. White arrowheads indicate the first detectable close-contact marked by accumulation of fluorescent CD58-HaloTag expressed by the U-2 OS cells. Images are displayed as maximal-intensity projections of the focal stack. Scale bars, 5 µm. **c**. Fraction of CD8^+^ T-cells forming close contacts with U-2 OS cells after 5’ (n=3 and n=2 videos for U-2 OS_ICAM-1_ and U-2 OS_WT_, each with >20 T cells). Error bars indicate SD. Data were compared using a Welch t test. **d-g**. Image-based analysis of the interactions of CD8^+^ T-cells with U-2 OS_WT_ and U-2 OS_ICAM-1_ cells (n=6 videos each with >20 interacting T cells). **d**. Average individual close contact area versus time. **e**. Average number of close contacts versus time. **f**. Average summed area of close contacts per T cell versus time. **g**. Average cumulative area of the target-cell surface scanned by single T-cells versus time. Imaging experiments were performed using CD8^+^ T-cells from at least two donors. Statistical comparisons, Fig. 3.

To analyse their structures and the dynamics of close-contact formation, we used custom-written image analysis software to segment the widefield images, building on our previous implementation of the method^9^. The CD58-marked close contacts, formed by T cells during interactions with both types of cells, had areas in the range of 0.1-0.2 µm^2^ throughout the interaction, as observed on SLBs^1,9,35^, with only a gradual increase for longer-lived interactions (Fig. 2d). For T cells interacting with U-2 OS_WT_ cells, the average number of close contacts slowly increased following initial contact, reaching a limit of ∼5 per cell (Fig. 2e). The presence of ICAM-1 dramatically altered the dynamics of close-contact formation. The average number of close contacts increased rapidly, peaking at ∼15 per cell (Fig. 2e). The combined area of close contact formation (Fig. 2f) and the cumulative area of target cell surface that was sampled (Fig. 2g) both increased dramatically in the presence of ICAM-1.

### Impact of TCR engagement on close-contact formation

To study the impact of TCR ligation on close-contact formation, N-Med/gp100-HLA-A2 complexes were presented at two densities on the surface of the U-2 OS cells, *i*.*e*., low (∼1 N-Med/µm^2^; ^Lo^U-2 OS_WT_ and ^Lo^U-2 OS_ICAM-1_ cells) and high (∼60 N-Med/µm^2^; ^Hi^U-2 OS_ICAM-1_ cells; Fig. S5a-d). Primary T-cells exhibited similar maximal responses, measured as CD69 and CD25 upregulation, to both U-2 OS_WT_ and U-2 OS_ICAM-1_ cells presenting TCR ligands at the two densities (Fig. S5e,f). The interactions of T cells with their targets were imaged using two-color epifluorescence widefield microscopy and eSPIM. For close-contact analysis, we used the interactions of T cells with ^Lo^U-2 OS_WT_ and ^Lo^U-2 OS_ICAM-1_ because the ligand density was a closer match to the physiological densities of native TCR ligands^36,37^. As was the case in the absence of ligand, the fraction of T cells forming close contacts, and the rate at which the contacts formed, was greatly accelerated by ICAM-1 (Fig. 3a-c; Fig. S5g,h; Supplementary Movies 5-8). The presence of ICAM-1 also enhanced the rate at which T cells killed the target cells (Fig. 3d) and increased ligand sensitivity of the T cells [6.4-fold and 4.1-fold decrease in the EC_50_(CD69) (Fig. 3e) and EC_50_(CD25) (Fig. S5i-k) values, respectively]. Surprisingly, the presence of the TCR ligand did not substantially alter close-contact dynamics: after 5 minutes of contact, the numbers of close contacts (Fig. 3f,g) and their total footprint (Fig. 3h,i) were unchanged at an average of 3 contacts and 0.4 µm^2^ per cell, and 13 contacts and 2.8 µm^2^ per cell for T cells interacting with ^Lo^U-2 OS_WT_ and ^Lo^U-2 OS_ICAM-1_ cells, respectively. Close-contact dimensions were also unaffected by the presence of the ligand: average close-contact areas (Fig. 3j) and diameters (Fig. 3k) were ∼0.2 µm^2^ and ∼500 nm, respectively, for T cells interacting with ^Lo^U-2 OS_WT_ and ^Lo^U-2 OS_ICAM-1_ cells, with the dimensions remained largely unchanged through the course of contact (Fig. 3l). The cumulative area of target cell surface that was sampled (Fig. 3m,n) was also unchanged. Overall, close-contact structure *per se* was unaffected by adhesion or stimulation. At later time points, the close contacts frequently displayed centripetal motion resulting in the formation of structures resembling multifocal synapses^38^, comprised of individual CD58 puncta (Fig. 3b; Supplementary Movie 6).

**Figure 3.**
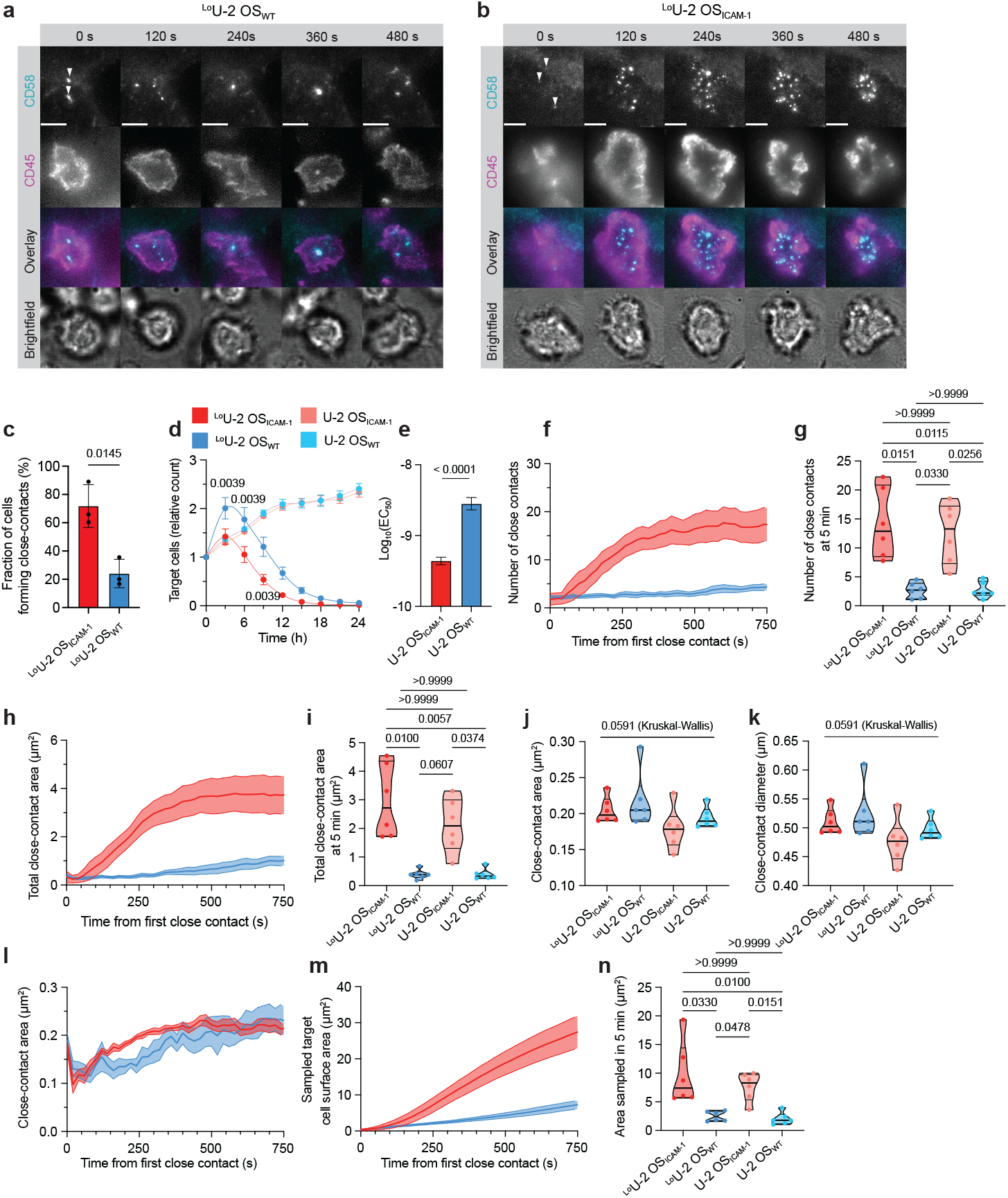
Efficient close-contact formation enhances T-cell activation and accelerates target-cell killing. **a, b**. Epifluorescence-based widefield imaging of CD8^+^ T-cells interacting with a monolayer of ^Lo^U-2 OS_WT_ (a) and ^Lo^U-2 OS_ICAM-1_ (b) cells presenting N-Med ImmTACs. White arrowheads indicate the first detectable close-contacts marked by accumulation of fluorescent CD58-HaloTag expressed by the U-2 OS cells. Images are displayed as maximum-intensity projections. Scale bars, 5 µm. **c**. Fraction of CD8^+^ T-cells forming close contacts with N-Med ImmTAC-presenting U-2 OS cells after 5’ (n=3 videos). Error bars show SD. Data were compared using a Welch t test. **d**. Killing of U-2 OS cells by CD8^+^ T-cells over time measured using Incucyte imaging. Killing assays were performed using CD8^+^ T-cells from three donors, in triplicate. Error bars show SEM. Data for ^Lo^U-2 OS_WT_ and ^Lo^U-2 OS_ICAM-1_ cells at selected time points (t = 3, 6, 9 h) were compared using a Wilcoxon signed-rank test. **e**. N-Med EC_50_ values measured in co-cultures of CD8^+^ T-cells with U-2 OS_WT_ and U-2 OS_ICAM-1_ cells. EC_50_ values were determined as in Fig. 1; see also Fig. S4. Error bars correspond to the 95% CI. EC_50_ values were compared using an F-test of normalised data. Three co-cultures were performed as biological repeats, with measurements in triplicate. Data were collected using CD8^+^ T-cells from at least two donors. **f-n**. Image-based analysis of CD8^+^ T-cells interacting with ^Lo^U-2 OS_WT_ and ^Lo^U-2 OS_ICAM-1_ cells [n=6 videos each with >20 interacting T cells; comparisons are made with T cells interacting with non ImmTAC-presenting U-2 OS cells (Fig. 2e-g)]. **f, g**. Average number of close contacts formed per T cell versus time (f) and 5’ after detecting the first close contact (g). **h, i**. Average summed area of close contacts per T cell versus time (h) and 5’ after detecting the first close contact (i). **j, k**. Average areas (j) and diameters (k) of all close contacts **l**. Average area of individual close contacts versus time. **m, n**. Average cumulative area of the target-cell surface scanned by a single T cell versus time (m) and 5’ after detecting the first close contact (n). Data were compared using a Kruskal-Wallis test and post-hoc Dunn’s test for multiple comparisons. Violin plots indicate median and quartiles. Imaging experiments were performed using CD8^+^ T-cells from at least two donors.

We also sought to confirm that the interactions of T cells with target cells presenting as model ligands were comparable to those with targets presenting conventional TCR ligands. For this, we generated primary CD8^+^ T-cells expressing the 1G4 TCR tagged at the C-terminus of the β-chain with a HaloTag (Fig. S5l, m). As targets, we used ^9V^U-2 OS_ICAM-1_ cells presenting the ‘9V’ variant of the cancer/testis NY-ESO-1 antigen (SLLMWITQV), which is presented by HLA-A2 and has a solution affinity of K_D_ = 7.2 µM for the 1G4 TCR^39^. The 1G4 TCR-expressing T-cells formed discrete, sub-micron sized contacts with the cells presenting 9V/HLA-A2, in the manner of T-cells interacting with N-Med-presenting cells and, likewise, did not form canonical, mono-focal synapses during the period of imaging (Fig. S5n, Supplementary Movie 9).

### Signaling at close contacts

To explore the spatiotemporal relationship of close-contact formation and signaling, we used spinning-disk confocal microscopy, which afforded better spatial resolution and optical sectioning than epifluorescence widefield microscopy or eSPIM. We first imaged N-Med accumulation at close contacts as a proxy of TCR engagement. Using the ICAM-1 presenting target cells permitted the analysis of many more productive interactions. The fluorescence signal from ^Lo^U-2 OS_ICAM-1_ cells (∼1 N-Med/µm^2^) was too weak to reliably track ligand accumulation, requiring the imaging of ^Hi^U-2 OS_ICAM-1_ cells (∼60 N-Med/µm^2^). Under these conditions, the N-Med ImmTACs readily accumulated within the close contacts, and the T-cells frequently formed conventional bulls-eye patterned synapses with CD58 and TCR accumulating centripetally (Fig. 4a,b; Supplementary Movie 10). To test whether close contacts are sites of TCR triggering and receptor phosphorylation, we generated primary CD8^+^ T-cells expressing Halo-tagged ZAP70 and Lck fusion proteins, each labelled with fluorescent JF646-HaloTag ligand (Fig. S6a,b). Cytosolic ZAP70 was recruited to sites of CD58 accumulation at the surface of ^Hi^U-2 OS_ICAM-1_ cells (enrichment: ∼1.38) with close-contact formation preceding ZAP70 accumulation (Fig. 4c-f; Fig. S6d; Supplementary Movie 11). In contrast, membrane-anchored Lck was typically more uniformly distributed (average enrichment: ∼1.09), with rare cells showing modest enrichment during the early stages of close-contact formation (Fig. 4e,f; Fig. S6c,e; Supplementary Movie 12). Despite the relatively high ligand density, not all close contacts produced similar levels of TCR phosphorylation (*i*.*e*., ZAP70 recruitment; Fig. 4e,f), and the variation could not be attributed to differences in the close-contact area (R^2^ 0.087; Fig. S6f). We also observed that ZAP70 recruitment occurs concomitantly with increases in cytosolic calcium levels (Fig. 4g-i). We previously showed that the formation of a single close-contact suffices to initiate signaling in response to cognate pMHC presented by SLBs^9^. This was also true for authentic target interactions: for most T cells interacting with ^Lo^U-2 OS_WT_ cells, a single close contact was detected at the time of calcium flux (Fig. 4j-l; Supplementary Movie 13). For interactions with ^Lo^U-2 OS_ICAM-1_ cells, on average, five close contacts formed by the time of calcium flux, likely owing to the accelerated rate of close-contact formation (Fig. 4k,l; Supplementary Movie 14). Nevertheless, a small fraction of T cells interacting with ^Lo^U-2 OS_ICAM-1_ (∼7%) fluxed calcium before two or more close contacts were established. These observations established that close contacts are *bona fide* sites of ligand recognition and TCR phosphorylation.

**Figure 4.**
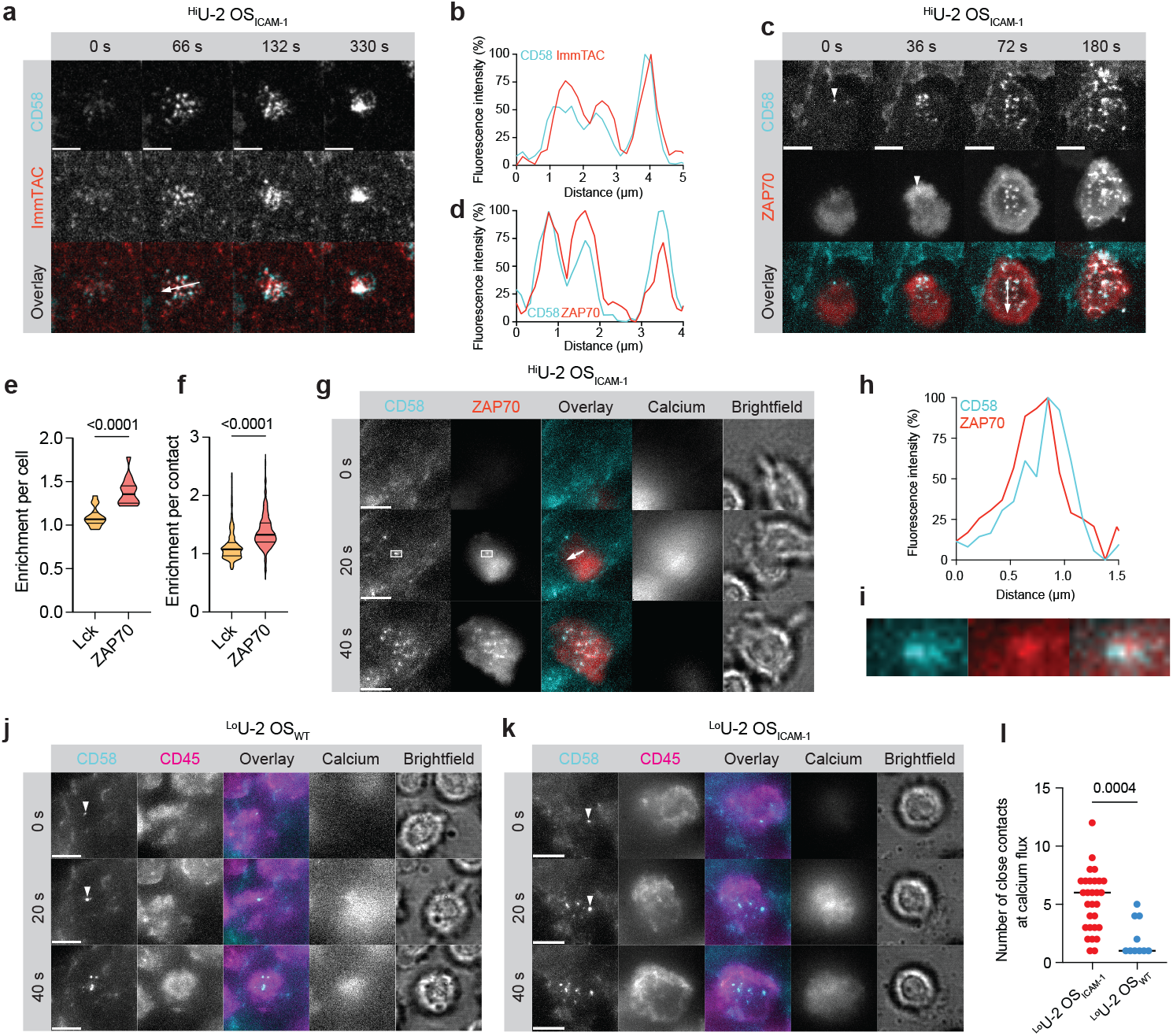
The TCR engages ligand and is triggered at close contacts. **a**. Spinning-disk confocal imaging of CD8^+^ T-cells interacting with ^Hi^U-2 OS_ICAM-1_ cells presenting fluorescent N-Med ImmTAC. **b**. Min/max normalised intensity line profile in the region of the white arrow in (a). **c**. Spinning-disk confocal imaging of ZAP70-HaloTag CD8^+^ T-cells interacting with ^Hi^U-2 OS_ICAM-1_ cells presenting N-Med ImmTAC. White arrowheads indicate the first detectable close contacts marked by accumulation of fluorescent CD58-HaloTag and recruitment of ZAP70-HaloTag. **d**. Min/max normalised intensity line profile in the region of the white arrow in (c). **e, f**. Enrichment of Lck-HaloTag and ZAP70-HaloTag at close contacts, calculated per cell (e) or per close contact (f). n=15 Lck-HaloTag-expressing and ZAP70-HaloTag-expressing T cells. Data were compared using a Mann-Whitney test. Violin plots indicate median and quartiles. **g**. Epifluorescence-based widefield imaging of ZAP70-HaloTag-expressing CD8^+^ T-cells labelled with a cytosolic calcium indicator Fluo-4 AM, interacting with ^Hi^U-2 OS_ICAM-1_ cells presenting N-Med ImmTAC. **h**. Min/max normalised intensity line profile in the region of the white arrow in (g). **i**. Close-up view of the close contact boxed in (g). **j, k**. Epifluorescence-based widefield imaging of CD8^+^ T-cells labelled with a cytosolic calcium indicator Fluo-4 AM interacting with ^Lo^U-2 OS_WT_ (j) and ^Lo^U-2 OS_ICAM-1_ (k) cells presenting N-Med ImmTAC. White arrowheads indicate the close contact marked by accumulation of fluorescent CD58-HaloTag detected prior to calcium flux. **l**. Number of close contacts formed by CD8^+^ T-cells interacting with ^Lo^U-2 OS_WT_ and ^Lo^U-2 OS_ICAM-1_ cells at the time of calcium flux (n=30 ^Lo^U-2 OS_ICAM-1_ cells, 10 ^Lo^U-2 OS_WT_ cells). Medians are indicated. Data were compared using a Mann-Whitney test. Imaging experiments were performed using CD8^+^ T-cells isolated from at least two donors. All images are displayed as maximal-intensity projections of the focal stack. Scale bars, 5 µm.

### CD45 exclusion

Numerous studies implicate CD45 segregation in the initiation of TCR signaling^1,9,11,13,16,17,35,40–44^, but it is yet to be directly observed during the early cellular interactions of T cells. To slow down the interaction dynamics in order to have the best opportunity to observe signaling-associated events, we collected data at 20°C. At the lower temperature, close-contact formation on ^Lo^U-2 OS_ICAM-1_ cells was slowed substantially (Fig. S7a-c), but the dimensions of the contacts were not significantly changed (Fig. S7e,f) and the contacts continue to recruit ZAP70 (Fig. S7g,h). In addition to labelling CD58 and CD45, we stained the cell membrane with CellMaskGreen to control for the effects of surface topography on the fluorescence signal; in this way, we observed that CD45 was depleted from close contacts (Fig. 5a-c). The average level of CD45 exclusion on a per-cell basis was 37% (*i*.*e*., enrichment: 0.63; Fig. 5d), and the maximum exclusion on a per-contact basis was 78% (enrichment: 0.22; Fig. 5e, Fig. S7i). Close-contact size and the degree of exclusion were uncorrelated (R^2^ = 0.046; Fig. S7j). As expected, cell membrane fluorescence was unchanged inside versus outside the close contacts (*i*.*e*., enrichment: 1.01; Fig. 5d,e). ZAP70 recruitment occurred simultaneously with CD45 exclusion at early close contacts (Fig. 5f,g).

**Figure 5.**
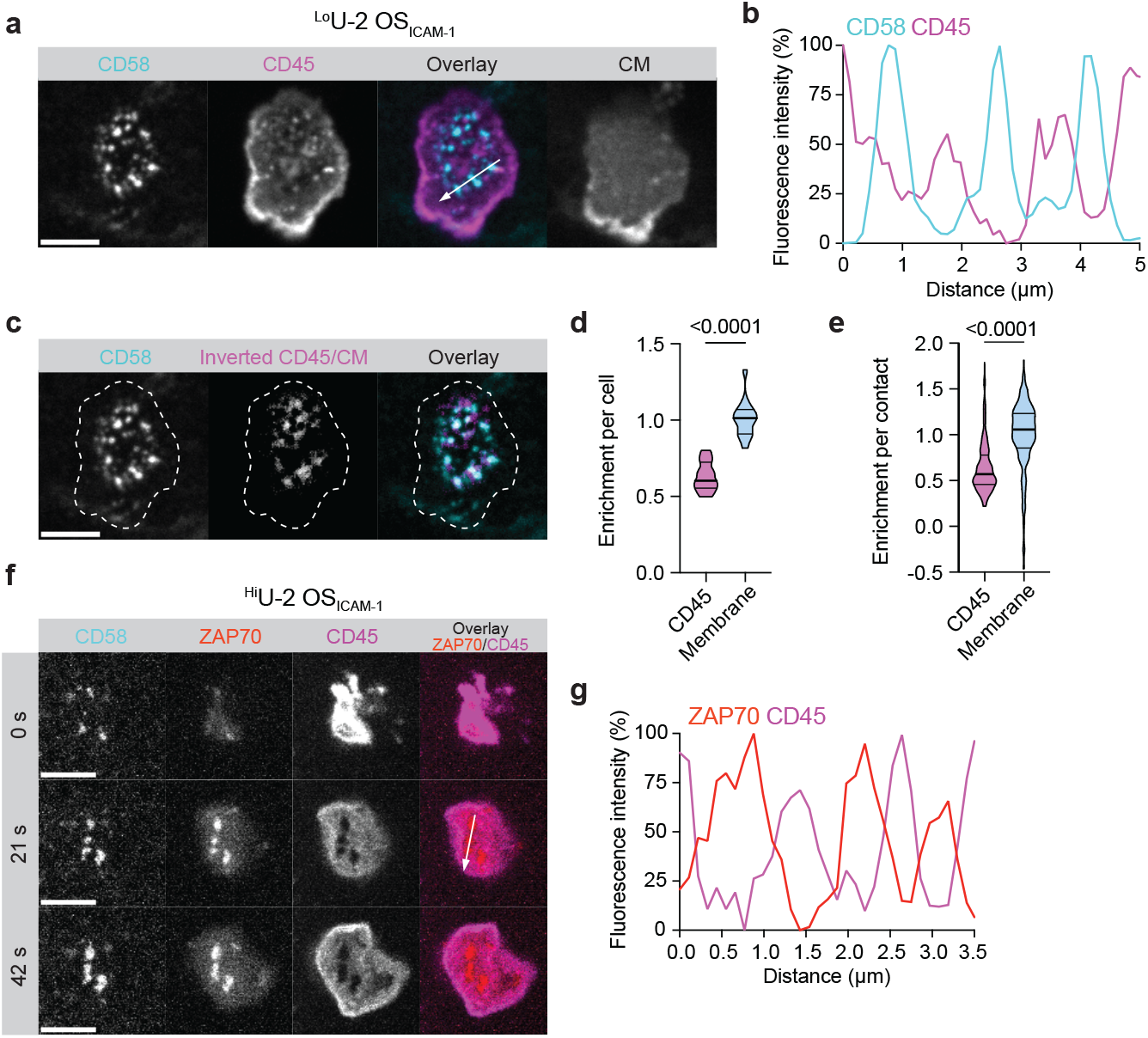
Partial depletion of CD45 at close contacts. **a**. Spinning-disk confocal imaging of CD8^+^ T-cells interacting with ^Lo^U-2 OS_ICAM-1_ cells presenting N-Med ImmTAC. CM = cell membrane **b**. Min/max normalised intensity line profile in the region of the white arrow in (a). **c**. Close contacts marked by accumulation of fluorescent CD58-HaloTag, and the inverted fluorescence signal for CD45 normalised against the cell membrane (producing bright areas marked by low CD45 intensity). A single focal plane corresponding to the T cell/U-2 OS cell interface is shown in the images. Scale bars, 5 µm. **d, e**. Analysis of CD45- and cell membrane-fluorescence enrichment within close contacts calculated per cell (d) or per close contact (e; n = 15 cells). Data were compared using a Wilcoxon signed-rank test. **f**. Spinning-disk confocal imaging of ZAP70-HaloTag CD8^+^ T-cells interacting with ^Hi^U-2 OS_ICAM-1_ cells presenting N-Med ImmTAC. Images are displayed as maximal-intensity projections of the focal stack corresponding to the T cell/U-2 OS cell interface. Scale bars, 5 µm **g**. Min/max normalised intensity line profile in the region of the white arrow in (f).

### Impact of partial CD45 exclusion on TCR-proximal signaling networks

We were interested to know to what extent T-cell signaling networks are perturbed by receptor triggering, since so few and such small contacts form, and the changes in kinase/phosphatase ratio accompanying contact formation are also limited. Previously, mass cytometry was used to follow early signaling events in T cells, but only under conditions of very strong activation, *i*.*e*., by cross-linking receptor-bound, biotinylated anti-CD3 and anti-CD28 antibodies with streptavidin^45^. We used the method to characterise T-cell signaling under more physiological conditions using 44 heavy metal-conjugated antibodies allowing signaling-effector phosphorylation and cell state changes to be detected (Table S3)^46,47^. We used, as targets, A375 melanoma cells, which are HLA-A2 positive and, when loaded with ImmTACs, induce very potent T-cell cytotoxicity^48,49^. Unexpectedly, UMAP analysis of the single-cell mass cytometric data revealed that the overall signaling fluxes induced by N-Med were very limited and largely indistinguishable from those in unstimulated cells (Fig. 6a). In contrast, the phosphatase inhibitor pervanadate produced very large effects. We calculated the Earth Mover’s Distance to quantify the post-translational modification intensities for each marker, confirming that the pervanadate-treated cells formed the distinct cluster owing to large changes in the pLck, pSrc, pZAP70, pSLP-76, and pERK signals (Fig. S8). Stimulation with PMA/ionomycin, which bypasses early signaling events, only induced changes in downstream signaling effectors (*e*.*g*., MAPKAP2, p90RSK, pCREB). For the pervanadate and PMA/ionomycin treatments, the differentially-phosphorylated proteins exhibited dynamic changes in phosphorylation with time, which was not the case for N-Med treated cells (Fig. 6b). Replicate N-Med- and non-stimulated cultures were kept for examining the functional effects of the treatments and this showed that, despite the small effects on signaling, the N-Med-presenting A375 cells induced high levels of both T-cell CD69 expression (Fig. 6c) and cytotoxicity (Fig. 6d). This suggests that modest changes in the local balance of kinase/phosphatase activity at just a few close contacts triggers strong T-cell responses, with minimal changes in measurable signaling fluxes.

**Figure 6.**
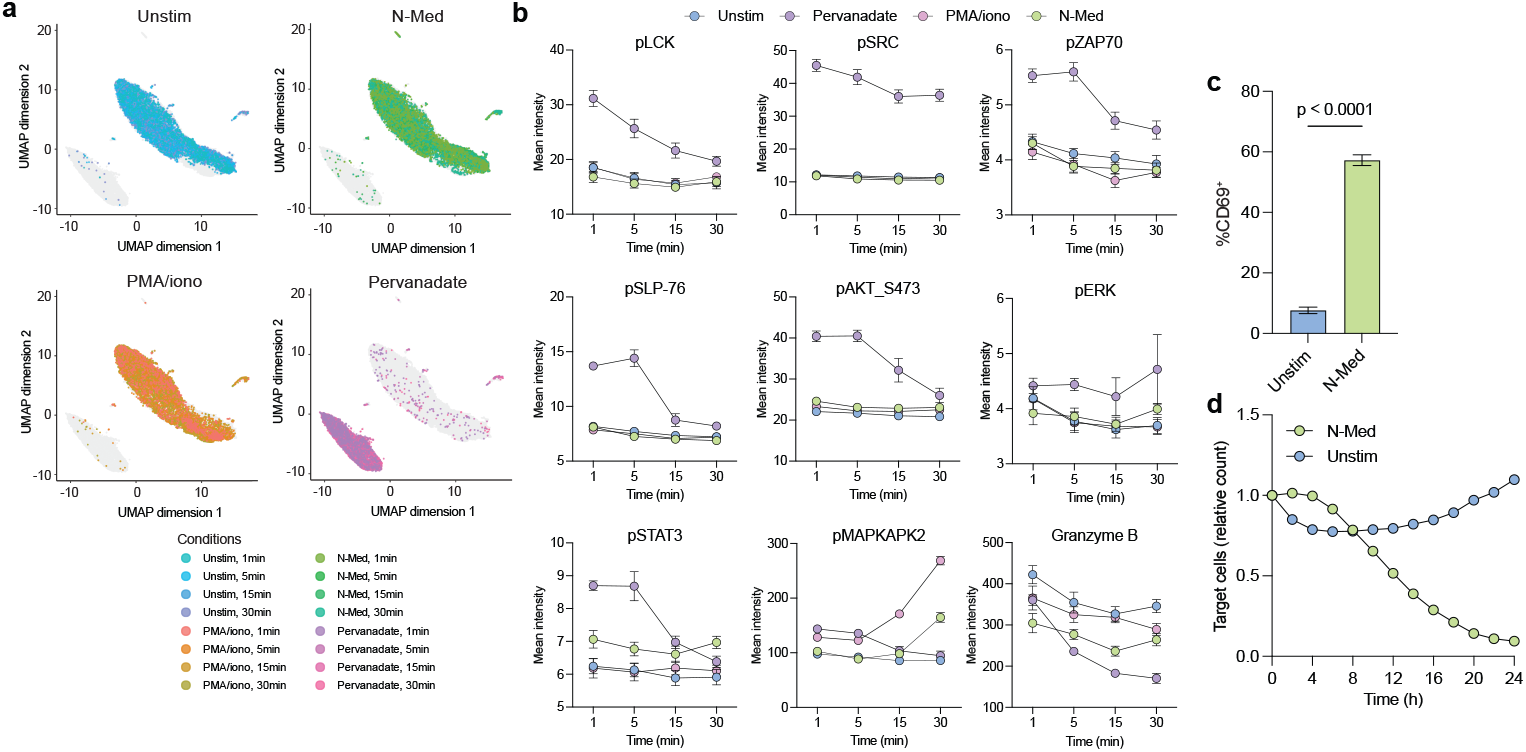
Changes in proximal signaling elicited by TCR triggering at close contacts. **a**. UMAP visualisation of signal intensities for CD8^+^ T cells isolated from two donors left unstimulated or stimulated with N-Med-presenting A375 cells, PMA/ionomycin, or pervanadate for 1, 5, 15, and 30 min. Intensities were measured by mass cytometry and analysed using CyGNAL. All cells are shown in grey, with overlays colored by treatment and timepoint. Four replicates for each treatment/timepoint condition. Co-cultures were performed as two biological repeats in duplicate. **b**. Levels of selected signaling effectors in (a). Error bars indicate SEM. **c**. Proportion of CD69^+^ T-cells following co-culture with N-Med-presenting A375 cells. n=3; error bars correspond to SEM. Co-culture assays were performed as three biological repeats in triplicate. Data were collected using CD8^+^ T-cells isolated from three donors. Data was compared using a Mann-Whitney test. **d**. Killing of A375 cells by CD8^+^ T-cells over time measured using Incucyte imaging. Killing assays were performed as three biological repeats in triplicate. Data were collected using CD8^+^ T-cells isolated from three donors.

### Stochastic simulations of TCR triggering at close contacts

At both authentic T-cell/target contacts and on advanced SLBs^9^, we find that CD45 is only partially (∼40%) excluded from close contacts. We conclude that this reflects the physicochemical properties of the contact interface^50,51^. However, it raises the question of whether the proximal signaling network is optimised for sensitive antigen detection at this level of local phosphatase depletion, as close contacts form. To explore this, we performed spatial stochastic simulations of TCR triggering at close contacts using Smoldyn^52^.

In our first model (Model 1, Fig. 7a and Fig. S9a) TCRs inside a close contact undergo sequential phosphorylation at a rate *k*_*phos*_ up to nP times (Fig. S9b), incorporating kinetic proofreading but without there being a strict requirement for ligand binding (the so-called “dwell-time” or KS-KP model)^53,54^. We allowed a single TCR reaching nP phosphorylations to initiate signaling and defined the true positive rate (TPR) as the fraction of simulations in which this triggering condition is met for a given ‘scan time’, at a ligand density of 10/µm^2^. The scan time corresponds to the lifetime of the close contact and sets an upper time limit to the triggering condition. The false positive rate (FPR) is determined from ligand-free simulations. Varying *k*_*phos*_ between 0.5-5 s^-1^, with a scan time of 7.5 s (ref. 1) ^1^we found that for nP = 5, the TPR exceeded the FPR at lower phosphorylation rates, whereas the FPR approached 1 at *k*_*phos*_ = 4 s^-1^ (Fig. 7b). To maximise the TPR while keeping the FPR low, the statistic to be optimised is TPR^*^(1-FPR), which is maximal at lower *k*_*phos*_ (Fig. 7c). Increasing the scan time shifted the optimal *k*_*phos*_ to lower values (Fig. 7d). For the optimal *k*_*phos*_ at a given scan time, increasing nP leads to an increase in *k*_*phos*_ (Fig. 7e and Fig. S9c). At the same time, a wider range of *k*_*phos*_ values resulted in higher values of the TPR^*^(1-FPR) statistic for nP = 7 compared to nP = 5 (Fig. S9d). The measured value of *k*_*phos*_ = 2.2 s^-1^ was the optimal value we observed for nP >= 4P at scan times below 7.5 s (Fig. 7e)^54^.

**Figure 7.**
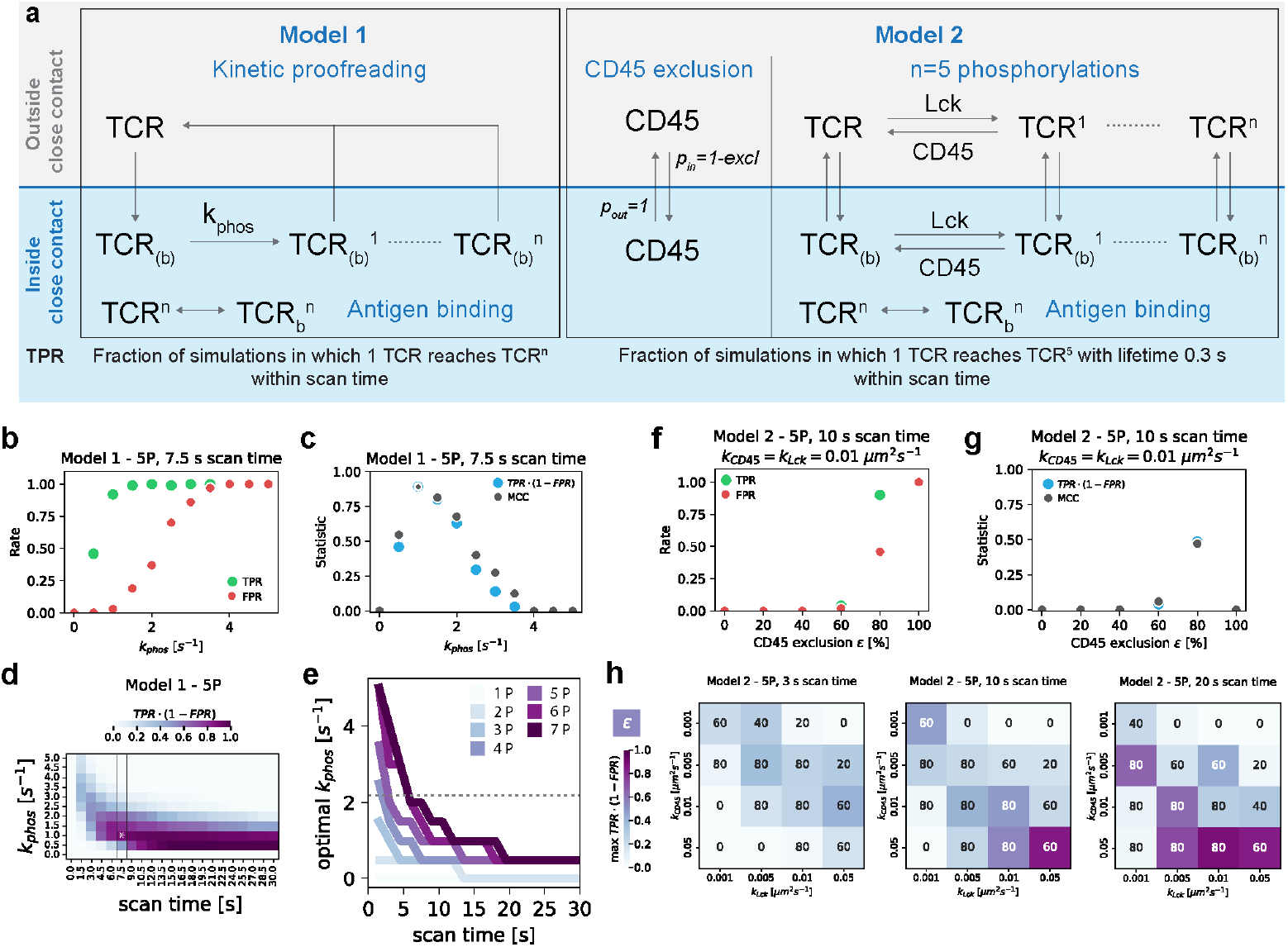
Partial exclusion of CD45 as a prerequisite for optimal antigen discrimination. **a**. Overview of Model 1 and 2. Reactions are separated into those occurring inside and outside the close contact (CC). In both models, the TCR can freely diffuse in and out of the CC, but antigen binding is only allowed inside the CC. In Model 1, TCR phosphorylation is described by a single forward rate, k_phos_, and is only possible inside the CC. In Model 2, TCR phosphorylation is modelled as forward and backward reactions both inside and outside of the CC. CD45 exclusion inside the CC is determined by the out-to-in transition probability p_in_. **b-e**. Results for Model 1 (n=100 simulations). **b**. True positive rate (TPR) and false positive rate (FPR) for varying k_phos_, for a triggering endpoint of 5 phosphorylations and 7.5 s scan time, calculated from 100 simulations. **c**. TPR^*^(1-FPR) value and Matthew’s correlation coefficient (MCC) as summary statistics for (b). We assume that the T-cell proximal signaling machinery is optimized to maximize these statistics for TCR triggering. **d**. Heatmap indicating the TPR^*^(1-FPR) value for 5 phosphorylation events varying k_phos_ and scan time. The box at a scan time of 7.5 s highlights the same plot as in (c), with the asterisk marking the highest summary statistic along it. For longer scan times, lower k_phos_ values lead to higher summary statistics. **e**. For varying scan time, the k_phos_ value with the highest summary statistic (asterisk in d) is plotted. Different lines indicate varying numbers of phosphorylations as triggering end points. The dotted line indicates the literature value k_phos_= 2.2 s^-1 46^. Along this line, the more phosphorylations that are required, the longer the optimal scan time. **f-h**. Results for Model 2 (n=50 simulations). **f**. TPR and FPR for varying CD45 exclusion, for a triggering endpoint of 5 phosphorylation events and 10 s scan time. **g**. TPR^*^(1-FPR) value and MCC as summary statistics for (f). **h**. Optimal exclusion value (inset number) and summary statistic (color) for the k_CD45_ and k_Lck_ parameter space. Three different scan times are shown.

To more explicitly study the influence of varying CD45 density inside the contact, we expanded upon Model 1 and implemented reversible TCR phosphorylation^55^ by simulating the phosphorylation and dephosphorylation of the TCR by Lck and CD45 as second-order reactions with rate constants *k*_*Lck*_ and *k_CD45_*, respectively (Model 2, Fig. 7a). This allowed us to explicitly tune the CD45 density inside the contact via its outside-to-inside transition probability (Fig. S9e) and analyse TCR phosphorylation kinetics (Fig.

S9f). TPR and FPR were calculated as for Model 1, with the addition of a minimum lifetime requirement of the nP phosphorylated TCR for the initiation of downstream signaling (Fig. S9g). We used nP = 5, approximating the number of proposed KP steps between ligand binding and ZAP70 recruitment^56^. For *k*_*Lck*_ = *k_CD45_*= 0.01 s^-1^ and a scan time of 10 s, we again observed a higher TPR than FPR at intermediate exclusion values (Fig. 7f), with the highest value of the TPR^*^(1-FPR) statistic being observed for CD45 exclusion of 80% (Fig. 7g). We then varied *k*_*Lck*_ and *k_CD45_*independently from 0.001 to 0.05 s^-1^ and examined scan times ranging from 3 to 20 s (Fig. 7h). Increasing the scan time from 7.5 to 20 s improved the TPR^*^(1-FPR) statistic across the parameter space, allowing the identification of conditions under which TPR exceeded FPR (Fig. 7h, Fig. S9h,i). The narrow range of favourable exclusion values might be a reflection of the ultrasensitivity reported for the TCR proximal signaling network^57,58^. These simulations indicate that complete exclusion of CD45 is not necessarily the optimal trade-off between true positives and false positives, as the numbers of false positives scale non-linearly with the amount of exclusion. A range of conditions allowed a high TPR while keeping the FPR < 0.2. We previously observed a FPR of 20% for calcium signaling by T cells on SLBs^9^ and note that an initially high FPR is acceptable for multi-stage search problems^59^.

## DISCUSSION

We previously studied the interactions of T cells with advanced SLBs presenting a 40 nm-deep glycocalyx alongside the ligands of small (CD2) and large (LFA-1) adhesion molecules, and null and agonistic pMHC^9^. We found that T cells punch through the glycocalyx, forming remarkably uniform 0.4 μm-diameter structures we called “close contacts”. Close contacts were the sites of TCR engagement and signaling, and constraints on their size underpinned ligand discrimination. Cai *et al*. had previously observed the formation of similar microvillar-based contacts when T cells interact with SLBs presenting fluorescent quantum dots^1^.

Whether insights obtained using these hard, planar bilayers were relevant to the interactions of T cells with softer, more topographically complex cellular targets, often presenting much deeper glycocalyces^18^, was unclear. In particular, it was uncertain whether size-constrained close contacts would form. Here, we studied the interactions of T cells with a tumor-derived cell line, U-2 OS, presenting a model TCR ligand, *i*.*e*., a bi-specific ImmTAC, and a ∼250 nm-deep glycocalyx. Having established that ImmTACs faithfully mimic the signaling activity of native, agonistic pMHC insofar as ImmTAC-induced signaling was sensitive to binding affinity, receptor/ligand complex dimensions, and co-receptor binding activity^12,25,30^, we showed that T cells formed uniform, size-constrained (0.5 μm) close contacts with tumor cell targets also. T cells interacting with tumor cells presenting native pMHC rather than ImmTACs formed contacts with the same structure and similar dynamics.

A key finding was that the rate of close-contact formation on U-2 OS cells was substantially lower than on SLBs, presumably due to the extra thickness of the tumor cell glycocalyx. Any heightened barrier activity of the tumor-cell glycocalyx was, however, readily overcome by interactions with ICAM-1. Strikingly, the fraction of T cells forming contacts increased 3-4 fold in the presence of ICAM-1, and the number of close contacts per cell and the rate of target-cell scanning were 5- and 4-fold higher, respectively. A presently unexplained difference between SLBs and the U-2 OS cells is that these effects of ICAM-1 were not TCR ligand-dependent, *i*.*e*., driven by “inside-out” signaling^9,60^. Nevertheless, despite the reduced target-scanning rate, non-ICAM-1-expressing tumor cells were completely eliminated from cultures by the primary T-cells with a delay of just 3h over a 24h period versus ICAM-1-expressing cells, underscoring the sensitivity and efficiency of T-cell killing. Even so, to eliminate tumors, T cells need to infiltrate malignant tissue and respond quickly, with only small fractions (∼10%) of the infiltrating T-cells typically engaging the tumors^61–64^. Therefore, target-scanning rate is likely to impact tumor control, and marked benefits could result from optimising the adhesiveness of therapeutic T-cells.

A large body of data suggests that the exclusion of CD45 is a pre-requisite of the TCR triggering mechanism ^5,11–17^. However, this had not been demonstrated at authentic T-cell/target cell contacts, undermining confidence in the kinetic-segregation model of TCR triggering^65^. Using advanced imaging, we confirmed that CD45 is excluded but only partially (37% on average). Partial exclusion was anticipated by some experiments with model systems, but it was important to confirm that it occurs at genuine cellular contacts and to measure the extent of exclusion, since experiments with the model systems produced mixed results. In lipid vesicle-based models, Schmid *et al*. observed that proteins 2 nm larger than a 10 nm membrane gap are fully excluded from adhesive contacts^6^, however Carbone *et al*. observed only 20% exclusion of CD45R0 (21.5 nm) and 40-50% exclusion of CD45RABC (>21.5 nm) at a 15 nm gap^66^. For cells interacting with bilayers, we observed 60% depletion of CD45 at close contacts formed by CD2 only^5^, and 30-40% exclusion on bilayers also presenting ICAM-1 and a glycocalyx^9^. Similarly, at cellular interfaces, Alakoskela *et al*., showed that spherical objects (*i*.*e*., quantum dots) as much as 10 nm larger than the adhesive gap are only 40-50% excluded^67^. Finally, we have shown that strongly agonistic antibodies are only 2-fold better at locally excluding phosphatases than weaker agonists^68^. The overall concordance of this data suggests that the extent of CD45 exclusion very likely reflects the physicochemical properties of the proteins and the interface, *i*.*e*., the sum of multiple factors, including entropic effects, the extent of allowable membrane deformation, molecular crowding, electrostatic repulsion, and the degree to which proteins can rotate about their anchoring points^6,69,70^.

That CD45 is only weakly excluded from close contacts suggests that receptor triggering in T-cells is sensitive to small, local changes in the ratio of kinase and phosphatase activities. Perhaps accordingly, but certainly also reflecting the small size and the relatively paucity of contacts that form, our mass cytometric analysis showed that T-cell activation relies on very limited changes in the global levels of phosphorylation of key downstream effectors of signaling, to the extent that these changes were largely undetectable using this method. Previous calculations have nevertheless suggested that partial CD45 exclusion suffices to initiate signaling ^54^, and our stochastic simulations also imply that the proximal signaling network could be optimised for discriminatory ligand detection in this setting. This is because the false-positive receptor triggering-rate scales with exclusion and, at the limit, signaling becomes ligand-independent (*i*.*e*., the FPR reaches a value of 1). We showed previously that TCR signaling can be triggered artificially by a small (5-fold) change in receptor diffusion-rate, also emphasizing the sensitivity of the triggering mechanism^17^.

Finally, and notably, we did not observe large, highly-structured “bulls-eye”-like immune synapses forming in the course of our imaging experiments under conditions of physiological ligand expression, when they would have been expected to form (*i*.*e*., within 20 minutes^71^). Occasionally, multi-focal accretions of close contacts formed and it’s possible that larger structures could appear later. Our data show instead that discriminatory signaling and the initiation of T-cell activation, at the very least, rely on fairly subtle changes in kinase/phosphatase activity, acting on small numbers of signaling effectors at multiple, tiny cellular contacts.

### Limitations of the study

We have demonstrated that CD45 is excluded from close contacts at the time of signaling when T cells interact with authentic cellular targets. We made averaged measurements of the level of exclusion, and it will be important now to understand how much the level of CD45 exclusion varies at individual contacts and the minimum exclusion level required to initiate signaling. Since we showed that calcium signaling can be triggered by the formation of a single contact, the relationship between the degree of CD45 segregation and signaling can now be explored. Our simulations focus on early TCR triggering events in single close contacts and provide an explanation for our observed CD45 exclusion data. While we explored a broad range of kinetic parameters in our reaction network, additional mechanisms such as feedback modulation of the activity of Lck or downstream signaling will likely also affect the cell-wide T-cell response. An additional limitation of the study is that it has focussed on a single tumor-derived cell line, and it is not clear how the glycocalyces of other tumor cells will alter the probability of contact formation or the structure of the close contacts that form. Finally, our experiments were undertaken in the two-dimensional setting of a tumor cell monolayer, and it’s uncertain how cell contact might differ in the crowded three-dimensional environment of tumors, where T cells undergo drastic morphological transitions in the course of locating their targets^61^.

## Acknowledgements

This work was funded by the Wellcome Trust (grant 207547/Z/17/Z to S.J.D.), CRUK (grant DRCCIPA\100010 to S.J.D. and D.K.), the MRC (grant MR/W025507/1 to E.P., S.S., C.T., and S.J.D.) and the Royal Society (Research Professorship RP150066 to D.K.). The authors thank Professor Omer Dushek (Oxford University) for helpful comments on the stochastic simulations.

## Author contributions

The experiments were conceived by SFL, AMS, DK, and SJD, and the methodology developed by MKot, DFH, MKör, DB, SS, JC, and CJT. The experimental work was undertaken by MKot, DFH, MKör, DB, AMS, ZZ, SK, JM, BL, JC, MF, GB, DKC, and BW. Data analysis was performed by MKot, DFH, MKör, ZZ, BF, YL, SFL, HC, SS, and EP. The work was supervised by SS, EP, CJT, SFL, AMS, DK, and SJD, and funding was acquired by SS, CJT, EP, DK, and SJD. The original manuscript was written by MKot, MKör, AMS, and SJD, and revised by all the authors.

## METHODS

### Cell culture

All cell culture was performed in HEPA-filtered cell culture cabinets. All media components were bought sterile or 0.22 μm filtered before use. Cells were grown at 37 °C in 5% CO_2_. Cell density and viability were monitored with a 1:1 mix of cell culture medium and 0.4% Trypan blue. Stained cells were analysed with a Countess II automated cell counter (ThermoFisher). Cells were independently tested negative for mycoplasma (Translational Immune Discovery Unit, WIMM). Primary CD8^+^ T cells were cultured in Roswell Park Memorial Institute 1640 Medium (RPMI 1640; ThermoFisher, Cat# 21875034) supplemented with 10% (v/v) fetal bovine serum (FBS; ThermoFisher, Cat# 10500064), 10 mM HEPES (ThermoFisher, Cat# 15630080), 50 U/ml Penicillin, 50 μg/ml Streptomycin, 100 μg/ml Neomycin (Sigma-Aldrich, Cat# P4083), 2mM L-Glutamine (Sigma-Aldrich, Cat# G7513), 1 mM Sodium pyruvate (Sigma-Aldrich, Cat# S8636), 50 U/ml human IL-2 (Biolegend, Cat# 589104). This medium is further referred to as complete RPMI. For the expansion of primary T cells using feeder cells, the concentration of human IL-2 was increased to 200 U/ml. U-2 OS cells were cultured in McCoy’s 5A medium (Sigma-Aldrich, Cat# M9309) supplemented with 10% (v/v) FCS, 50 U/ml Penicillin, 50 μg/ml Streptomycin, 100 μg/ml Neomycin. This medium is further referred to as complete McCoy’s. mOrange A375 cells generated as previously described^73^ and were cultured in Dulbecco’s Modified Eagle Medium/Nutrient Mixture F-12 (DMEM/F12; ThermoFisher, Cat# 11320033) supplemented with 10% (v/v) FBS, 50 U/ml Penicillin, 50 μg/ml Streptomycin, 100 μg/ml Neomycin, 1 mM Sodium pyruvate and 10 mM HEPES. HEK 293T cells were cultured in Dulbecco’s Modified Eagle Medium (DMEM; ThermoFisher, Cat# 41965039) supplemented with 10% (v/v) FBS, 50 U/ml Penicillin, 50 μg/ml Streptomycin, 100 μg/ml Neomycin, 2 mM L-Glutamine, 1 mM Sodium pyruvate. This medium is further referred to as complete DMEM. For passaging, adherent cells were detached from the flasks using 5 mM EDTA (ThermoFisher, Cat# 15575020) solution in phosphate-buffered saline (PBS; ThermoFisher Cat# 10209252).

### Lentivirus production

1×10^6^ HEK 293T cells in 2 ml of complete DMEM were seeded onto 6-well plates 24 h prior to the transfection. 0.5 μg lentiviral envelope plasmid (pMDG, Addgene 187440), 0.5 μg packaging plasmid (p8.91, Addgene 187441) and 1 μg transfer plasmid (pHR or pHRi, Addgene 187358) were combined in 100 μl of unsupplemented DMEM medium. For the co-transfection, 4.5 μl of GeneJuice (Sigma-Aldrich, Cat# 70967) was added to the plasmid mixture and gently resuspended, following a 30-minute incubation at 25 °C the complexed DNA was added to the HEK293T cells. 48 to 72 h later, the supernatant containing the lentivirus was harvested and filtered (0.22 μm, PES). The lentivirus-containing supernatant was used immediately or stored at -80 °C for further use.

### Creation of U-2 OS cell lines

U-2 OS cell lines expressing variants of the gp100 single-chain trimer (SCT) were generated by lentiviral transductions. To generate U-2 OS_WT_, U-2 OS_ICAM-1_ and ^Hi^U-2 OS_ICAM-1_ cell lines, endogenous CD58 was deleted. Briefly, U-2 OS cells (ATCC) were electroporated with Cas9/gRNA ribonucleoprotein (RNP). Guide RNA (gRNA) was prepared by mixing 6.25 μl of 400 μM tracrRNA (IDT, Cat# 1072532) and 400 μM CD58-targeting crRNA (IDT, target sequence: GTACTCATGGGATTGTCCTA). The mixture of tracrRNA and crRNA RNA was then incubated at 95 °C for 5 min and cooled to 25 °C. To generate RNPs, the gRNA was combined with 2.5 μl of 10 μg/μl S.p HiFi Cas9 Nuclease V3 (IDT, Cat# 1081060) and incubated for 30 min at 25 °C. The expression levels of CD58 were assessed by flow cytometry 9 days after transduction, and the CD58-knockout cells were isolated by FACS. To generate U-2 OS_WT_, U-2 OS_ICAM-1_ and ^Hi^U-2 OS_ICAM-1_ cell lines, expression of CD58-HaloTag, truncated ICAM-1 and wild-type gp100-SCT was sequentially established in the CD58-knockout U-2 OS cells via lentiviral transductions. Expression of mOrange in U-2 OS_WT_ and U-2 OS_ICAM-1_ cell lines was established via lentiviral transduction. All lentiviral transductions of U-2 OS cells were performed by seeding 0.5×10^6^ U-2 OS cells in a 6-well plate in 1 ml of complete McCoy’s media and 1 ml of lentiviral supernatant. The expression levels of SCTs, CD58-HaloTag, truncated ICAM-1 and mOrange were assessed by flow cytometry 72 h after transduction, and the cell lines were sorted by FACS to achieve optimal surface expression.

### Primary CD8^+^ T-cell isolation and culture

T cells were isolated from blood leukocyte cones provided by the NHS Blood and Transplant unit, John Radcliffe Hospital, Oxford, UK. Blood cones were used under the ethical guidance of NHS Blood and Transplant. Peripheral blood mononuclear cells (PBMC) were isolated by Ficoll-Paque density gradient centrifugation using Lymphoprep (StemCell Technologies, Cat# 18061) according to the manufacturer’s protocol. Primary CD8^+^ T cells were isolated from PBMCs using a MACS system comprising CD8^+^ T Cell Isolation Kit (Miltenyi Biotec, Cat# 130096495), LS columns (Miltenyi Biotec, Cat# 130042401), and a manual magnetic separator (Miltenyi Biotec, Cat# 130091051) which were used according to the manufacturer’s protocol. 10^6^ purified primary CD8^+^ T cells were resuspended in 1 ml of complete RPMI media and plated in a 24-well plate (Corning) with 75 μl of αCD3/αCD28 Dynabead suspension (ThermoFisher, Cat# 11131D) per well (1:3 cell-to-bead ratio) and cultured for 4-5 days. Next, αCD3/αCD28 Dynabeads were removed, and the primary CD8^+^ T cells were pooled and transferred to T-25 flasks. Expanding primary CD8^+^ T cells were kept in culture for another 10 days and passaged every other day to maintain a density of 0.5-2×10^6^ cells per ml. On the final day, the cells were frozen and stored in LN_2_ until later use.

### Generation of 1G4αβ-HaloTag, ZAP70-HaloTag and Lck-HaloTag primary CD8^**+**^T-cells

250 µl of Retronectin (Takara Bio, Cat# T100B) solution at 25 μg/ml in PBS was used to coat non-treated 48-well plates (Corning, Cat# 351178) for 24-48 h at 4 °C. Next, the retronectin solution was removed and plates were blocked with 300 μl of a 2% (w/v) BSA solution in PBS for 30 min at 25 °C. Next, the BSA solution was removed, and the wells were washed with 1 ml of PBS. 1.5 ml of the lentiviral supernatant was added to each well and the plate was centrifuged at 2000 x *g* for 90 min at 32 °C. PBMCs were isolated as previously described and plated at 2×10^6^ cell in 2 ml of complete RPMI media with 50 μl of αCD3/αCD28 Dynabead suspension (ThermoFisher, Cat# 11131D) per well (1:1 cell-to-bead ratio). 2 days later, 10^6^ cells, predominantly a mix of CD4^+^ and CD8^+^ T cells, were seeded in 1 ml of complete RPMI per well in the lentivirus-coated 48-well plates and the plate was centrifuged at 500 x *g* for 1 min at 25 °C. T cells were cultured for 4 days and passaged every other day. Transgene-positive CD8^+^ T cells were isolated using fluorescence-activated cell sorting. T cells transduced with the polycistronic 1G4α-1G4βHaloTag-LNGFR constructs were co-stained with an PE 9V-HLA-A2-streptavidin teteramer (Biolegend, Cat# 405204), FITC anti-NGFR (Biolegend, Cat# 345103) and AF647 anti-CD8 (Biolegend, Cat# 344725) antibodies. T cells transduced with the ZAP70-HaloTag and Lck-HaloTag constructs were co-stained using Janelia Fluor 646 HaloTag ligand (Promega, Cat# GA1120) and Pacific Blue anti-CD8 antibody (Biolegend, Cat# 344717). 0.5-1×10^5^ T cells were then seeded in a T-25 flask with 8 ml of feeder cells prepared 24 h earlier. 5 days later, 5 ml of complete RPMI media was added to the flask without disturbing the proliferating T cells. T cells were further cultured in a T-25 flask until a confluent layer of T cells at the bottom of the flask became visible. Next, 20 ml of complete RPMI media was added, and T cells were transferred into a T-75 flask. 14 days after FACS, T cells were frozen and kept in LN_2_ until further use.

### Preparation of feeder cells

PBMCs, from at least three donors, were isolated and irradiated for 10 min at 3000 rad. The irradiated PBMCs were centrifuged at 300 x *g* for 5 min and resuspended in complete RPMI media at 2×10^6^ cell/ml. PBMCs from different donors were combined at equal volumes and supplemented with phytohemagglutinin-L (ThermoFisher, Cat# 00497703) at 2.5 μg/ml and anti-CD3ε antibody (Biolegend, Cat# 317326) at 50 ng/ml.

### Protein purification

CD3ε and CD3γ chains were individually expressed in BL21 *E. coli*. The inclusion bodies were dissolved in a 20 mL denaturation buffer (6M guanidine, 50 mM Tris pH 8.1, 100 mM NaCl, 10 mM EDTA, 20 mM DTT) and incubated at 37 °C for 30 min. The denatured protein was then added dropwise to a constantly stirred 1 L refolding buffer (4 M Urea, 0.4 M L-arginine, 100 mM Tris, pH 8.1, 2 mM EDTA, 1.9 mM Cystamine, 6.5 mM Cysteamine). The refolding buffer was stirred for further 10 min and dialyzed for 16 h against deionized water, and twice against 10 mM Tris buffer (pH 8.1). Monomeric CD3εγ heterodimer was purified using anion exchange and size exclusion chromatography. The protein was enzymatically biotinylated with the BirA enzyme.

gp100-HLA-A2-His_12_ and 9V-HLA-A2-AviTag was produced as previously described^74^. Briefly, HLA-A2-His_12_ and beta-2-microglobulin (β_2_M) were expressed in BL21 *E. coli*. HLA-A2-His_12_, HLA-A2-AviTag and β_2_M were purified from inclusion bodies and refolded in the presence of the gp100 peptide (YLEPGPVTV, GenScript) or 9V peptide (SLLMWITQV, GenScript). Monomeric pMHC was purified using size exclusion chromatography. 9V-HLA-A2-AviTag was enzymatically biotinylated with the BirA enzyme and combined with PE-conjugated streptavidin at a 1:5 ratio (Biolegend, Cat# 405204) to produce tetramers.

Production of soluble ImmTACs has been previously described^20^. Briefly, TCR α-chain and β-chain, fused to anti-CD3 scFv via GGGGS-linker, were cloned into bacterial expression vectors. Polypeptides were expressed in E. coli and were refolded from denatured inclusion bodies as soluble disulfide-linked heterodimeric ImmTACs and purified by anion exchange and size exclusion chromatography. In certain instances, TCR chains and fusions were cloned into mammalian expression vectors, and ImmTACs were produced using the Expi293 GnTI-Expression System (ThermoFisher). ImmTACs were purified using ProteinL-affinity and size-exclusion chromatography. The purity of all proteins was assessed by reducing and non-reducing SDS-PAGE and concentrations were measured using absorption at 280 nm (Nanodrop One, ThermoFisher).

### Fluorescent dye conjugation

Gap8.3 antibody and ImmTACs were labelled via random lysine labelling using the NHS ester forms of the relevant fluorescent dyes. For labelling of proteins with either silicon rhodamine (Spirochrome) or Janelia Fluor 646 (Tocris Bioscience, Cat# 6148), the dyes were purchased in a powder form and reconstituted in anhydrous DMSO (Sigma-Aldrich). The relevant NHS ester dye was added to the protein in 0.1 M sodium bicarbonate buffer to achieve a protein:dye ratio of 1:20 (Gap8.3 antibody) and 1:30 (ImmTACs). The reaction was allowed to proceed for at least 1 h at 20 °C. Unreacted free dye was then removed via size exclusion chromatography columns (Bio-Rad). For labelling of protein with Alexa Fluor dyes, Antibody Labelling Kit (ThermoFisher, Cat# 10799764, Cat# 10257062) was used according to the manufacturer’s protocol. The labelling of the protein was then confirmed via UV-vis spectrophotometry (Nanodrop One, ThermoFisher).

### Flow cytometry

0.5×10^6^ cells were counted, washed in PBS (0.01% NaN3; PBS azide), and then stained at 4 °C for 1 h using relevant antibodies or ImmTACs. The centrifugation between the wash steps was carried out at 757 x *g* for 3 min at 4 °C. Cells were washed twice in PBS azide, fixed in 1% (v/v) paraformaldehyde (PFA) in PBS. Samples were analysed using the Attune NxT (Life Technologies) flow cytometer unless specified otherwise. Compensation was performed where required. Data were analysed using FlowJo. All cell sorting was performed by the Weatherall Institute of Molecular Medicine FACS Facility.

### Quantification of molecule copy numbers on U-2 OS cell surface

U-2 OS cells were individually stained using AF647-conjugated ImmTAC (10 nM), PE anti-CD58 (1:100; Biolegend, Cat# 330928) or PE anti-ICAM-1 (1:100; Biolegend, Cat# 353106) antibodies. To quantify the copy number of surface molecules, either BD Biosciences Quantibrite PE calibration beads (BD Biosciences, Cat# 340495) or Quantum Alexa Fluor 647 MESF beads (Bangs Laboratories, Cat# 647A) were used according to the manufacturer’s protocol.

### Flow cytometry-based activation assays

2.5×10^4^ U-2 OS cells were plated in a 96-well bottom plate and cultured in complete McCoy’s media Optionally, the media were supplemented with gp100 peptide. 18 h later, the U-2 OS cells were washed twice with PBS. For co-culture assays with gp100-pulsed U-2 OS cells, U-2 OS cells were labelled with ImmTAC in complete RPMI media for 30 min at 37 °C, washed twice with PBS, and 5×10^4^ T cells in complete RPMI were added on top of the target cells. For co-culture assays of T cells with gp100-SCT U-2 OS cells, 5×10^4^ T cells in complete RPMI were added on top of the gp100-SCT U-2 OS cells and ImmTAC dilution in complete RPMI media. 18 h later, T cells were transferred from the 96-well flat-bottom plate to a 96-well U-bottom plate for staining. T cells were washed with ice-cold PBS and stained with Zombie Yellow Fixable Viability stain (Biolegend, Cat# 423104, 1:300) for 20 min at 25 °C. Next, T cells were washed with ice-cold PBS and stained with the antibody cocktail containing FITC anti-CD3 antibody (Biolegend, Cat# 300440, 1:200), PE-Cy7 anti-CD8 antibody (Biolegend, Cat# 344712, 1:200), AF647 anti-CD25 antibody (Biolegend, Cat# 302618, 1:200) and PB anti-CD69 (Biolegend, Cat# 310920, 1:200) prepared in ice-cold PBS for 45 min at 4 °C. Next, T cells were washed twice with ice-cold PBS and fixed in 1% (v/v) formaldehyde in PBS and stored at 4 °C until analysed using Attune NxT (Life Technologies) flow cytometer.

### Cytotoxicity assays

5×10^4^ U-2 OS cells expressing mOrange were plated in a flat-bottom 96-well plate in 200 µl of complete McCoy’s media with gp100 peptide at 10 µM and cultured in a humidified incubator at 37 °C and 5% CO_2_ for 18 h. Following the incubation, the media was removed, and the U-2 OS cells were washed with 200 µl of PBS. Next, U-2 OS cells were labelled with 10 nM ImmTAC (N-Med) solution in complete RPMI for 20 min and washed twice with 200 µl of PBS. No ImmTAC was added to the control wells. 1×10^5^ primary CD8^+^ T cells in 200 µl of complete RPMI media were added to each well on top of U-2 OS cells. Plates were immediately placed in an Incucyte live-cell imaging system (Sartorius) with a scan every 3 h. The number of U-2 OS cells was assessed using mOrange fluorescence signal and normalised to the first scan. Data analysis and processing was carried out using Incucyte GUI software (Sartorius).

### Analysis of signalling pathways using mass cytometry

A375 cells were plated at 5×10^4^ cells per well in 96-well flat-bottomed plates and pulsed with gp100 peptide at 10 µM for minimum of 4 hours. The A375 cells were washed once with PBS before addition of N-Med ImmTAC to a final concentration of 0.1 nM. N-Med ImmTAC was allowed to the gp100-pulsed A375 cells at 37 °C for a minimum of 30 minutes before the start of the experiment.

CD8^+^ T cells isolated from two donors were then spun onto the peptide-pulsed A375 cells presenting ImmTAC (500g, 1 min) to initiate adhesion and start the activation. At the end of the time course, the cells were removed from the plate to 96-well U-bottomed plates, and then pelleted in ice-cold PBS (1800 rpm, 3 min) before fixation in 100 μl of 1.6% PFA. Cells were fixed at 4 °C for a minimum of 20 minutes, before three washes in 200 μl of PBS. For comparison, the same CD8^+^ T cells were stimulated with PMA/ionomycin, or pervanadate, a pharmacological protein tyrosine phosphatase inhibitor. Mass cytometric analysis was undertaken as described previously^46^. Briefly, dead cells were stained using 0.25 mM 194 Cisplatin (Standard BioTools, Cat# 201194) and then the cells were incubated with thiol-reactive barcodes overnight at 4 °C. Unbound barcodes were quenched in 2 mM GSH and the cells then pooled. The cells were dissociated into single cells using 0.875 mg/ml Dispase II (ThermoFisher, Cat# 17105041), 0.2 mg/ml Collagenase IV (ThermoFisher, Cat# 17104019), and 0.2 mg/ml DNase I (Sigma-Aldrich, Cat# DN25) in C-Tubes (Miltenyi Biotec, Cat# 130096334) using a gentle MACS Octo Dissociator with Heaters (Miltenyi Biotec, Cat# 130096427, SCR_020271). The single cells were washed in cell staining buffer (CSB; Standard BioTools, Cat# 201068) and then stained with extracellular rare-earth metal conjugated antibodies for 30 min at room temperature. The cells were then permeabilised in 0.1% (v/v) Triton X-100/PBS (Sigma-Aldrich, Cat# T8787), 50% methanol/PBS (Fisher Scientific, Cat# 10675112), and stained with intracellular rare-earth metal conjugated antibodies for 30 min at room temperature. The cells were washed in CSB and antibodies were cross-linked to the cells using 1.6% (v/v) FA/PBS for 10 min. Finally, the cells were incubated in 125 nM 191 Ir/193 Ir DNA intercalator overnight at 4 °C before being washed, resuspended in cell acquisition solution plus (CAS+; Standard BioTools, Cat# 201244) with 2 mM EDTA (Sigma-Aldrich, Cat# 03690), and analysed using a CyTOF XT (Standard BioTools, Cat# SCR_02634) at 200-400 events per second.

Signal intensities were measured using CyTOF at 1, 5, 15, and 30 minutes post-stimulation with a panel of 44 markers. Eight control markers were excluded from downstream analysis. Raw data were preprocessed using FlowJo, and analysed using the CyGNAL package with default settings (github.com/TAPE-Lab/CyGNAL)^47^. To measure the similarity between different treatment/timepoint conditions, Earth Mover’s Distance (EMD) scores were calculated and normalized against the entire dataset.

### A375 CD8^+^ T-cell killing and cytotoxicity assays

For the A375 killing assays and CD69 expression analyses, CD8^+^ T-cells were co-cultured with mOrange A375 cells presenting N-Med ImmTAC as described for the mass cytometry experiments. A375 cell growth was then monitored using Incucyte live-cell imaging system (Sartorius), acquiring scans once every 2 hours for a period of 24 hours. The number of A375 cells was assessed using mOrange fluorescence signal and normalised to the first scan. Data analysis and processing was carried out using Incucyte GUI software (Sartorius). After 24 hours, the CD8^+^ T-cells were removed from co-cultures and stained for 20 minutes at 4 °C in 100 µl with the following antibody cocktail prepared in PBS: CD8 PeCy5 (1:200; Biolegend, Cat# 344769), CD69 BV785 (1:200; Biolegend, Cat# 310931), ef780 Live-Dead (1:1000, eBioscience, Cat# 65086514). Next, T cells were washed twice in PBS azide, before resuspension in 200 ul of 1% (v/v) formaldehyde in PBS and analysed using a BD LSR Fortessa X-20 flow cytometer.

### Surface plasmon resonance-based binding assays

The binding of ImmTACs to their ligands gp100-HLA-A2-His_12_ and biotinylated CD3εγ was assessed using surface plasmon resonance (SPR) as implemented by Biacore 8K (Cytiva). All measurements were performed at 37 °C. HBS-P+ or HBS-EP+ (Cytiva, Cat# BR100671, Cat# BR100669) buffers were used to prepare the serial dilutions of ImmTAC for measurements of binding to gp100-HLA-A2-His_12_ and biotinylated CD3εγ, respectively. All runs were reference subtracted as well as blank subtracted. Both buffers were filtered and degassed using a vacuum filtration system (0.22 μm, PES). For affinity and kinetic measurements of ImmTACs to gp100-HLA-A2-His_12_, Series S Sensor Chip NTA (Cytiva, Cat# 28994951) was used according to the manufacturer’s instructions and multi-cycle analysis was conducted. The gp100-HLA-A2-His_12_ (at 50 nM) was captured on the chip for 60 s. ImmTACs were injected for 60 s, and the dissociation was monitored for 900 s. A twofold serial dilution ranging from 200 nM to 1.56 nM was used. HBS-P+ was used as a running buffer. The NTA Chip was regenerated using the regeneration solution (350 mM EDTA in the HBS-P+ buffer) according to the manufacturer’s instructions. For affinity and kinetic measurements of ImmTACs to the biotinylated CD3εγ, Series S Sensor CAPture Chip (Cytiva, Cat# 28920234) was used, and a single-cycle analysis was conducted. The CAP reagent (Cytiva, Cat# 29423383) was injected for 300 s. The biotinylated CD3εγ (at 50 nM) was captured on the chip for 180 s. ImmTACs were injected for 120 s and the dissociation was monitored for 60 s before the next injection. A twofold serial dilution ranging from 800 nM to 6.25 nM was used. HBS-EP+ was used as a running buffer. The CAPture Chip was regenerated with the regeneration solution (6 M guanidine-HCl, 0.25 M NaOH) according to the manufacturer’s instructions. The analysis of binding kinetics was carried out using Biacore Insight Evaluation Software (Cytiva).

### Confocal imaging of U-2 OS cells for cell-surface calculations

10^4^ U-2 OS cells in 300 μl of complete McCoy’s media were seeded in a glass-bottom 8-chamber coverslip (Ibidi, Cat# 80807). The next day, U-2 OS cells were incubated with 5 μg/ml CellMask Deep Red Plasma Membrane Stain (ThermoFisher, Cat# C10046) in complete McCoy’s media for 5 min (37 °C, 5% CO_2_). Next, U-2 OS cells were washed three times with 300 μl of complete McCoy’s media and fixed 0.8% paraformaldehyde (ThermoFisher, Cat# 28906) and 0.2% glutaraldehyde (Sigma-Aldrich, Cat# G5882) in PBS for 30 min at 25 °C. Fixed U-2 OS cells were washed three times with 300 μl of PBS and imaged in PBS using Zeiss LSM 980 inverted microscope in the AiryScan mode. 63x, NA 1.4 oil-immersion objective was used and the sample was illuminated using a 633 nm laser (He-Ne). The pinhole size was set to 1 AU. The z-stacks were taken from the apical to the basal plasma membrane with an interplane interval of 200 nm. Laser power and detector settings, including gain and digital offset, were set to prevent signal saturation while maintaining dynamic range. Image acquisition was performed using Zeiss ZEN software. The surface area of U-2 OS cells was calculated using ImageJ. Binary mask for each of the planes, corresponding to the cross-section of the cells, was produced as follows. Images were thresholded using the Otsu algorithm at the midplane, particles were dilated (5x) and holes were filled to create contiguous masks. The circumference of each binary mask, except for the one corresponding to the apical plane, was multiplied by 200 nm (interplane interval). The resultant value was added to the area of the binary masks corresponding to the apical and the basal planes to produce an estimate of the U-2 OS cell surface area.

### Confocal imaging of U-2 OS cells for glycocalyx measurements

U-2 OS_WT++_cells in suspension were fixed using a solution containing 0.8% paraformaldehyde (ThermoFisher, Cat# 28906) and 0.2% glutaraldehyde (Sigma-Aldrich, Ca# G5882) in PBS for 20 minutes at 25 °C. After fixation, cells were washed three times with PBS. To label the glycocalyx and plasma membrane, fixed cells were incubated with Wheat Germ Agglutinin (WGA; 6.1 μM)-Star Red conjugate and JF549 (200 nM), respectively. Next, U-2 OS cells were washed three times with PBS and allowed to settle onto glass-bottom 8-chamber coverslip (Ibidi, Cat# 80807) for 60 minutes. Imaging was performed using a in spinning disk confocal microscope (details below). For the imaging of the cellular glycocalyx, spinning disk microscope (3i intelligent, see details below) was used to take z-stacks at 0.5 µm intervals, such that the full volume of the U-2 OS cells was captured. Analysis was performed using MATLAB. Briefly, for each 2D plate a line profile was drawn through the cell’s centre of mass and rotated in 1° increments over 180°, generating a continuous radial sampling of the entire 2D image plane. Gaussian profiles were fit to the fluorescence signal of WGA and CD58-HaloTag and used to calculate σ and FWHM values.

### Scanning electron microscopy of fixed primary CD8^+^ T-cells

13 mm glass slides were placed in 12-well plates and incubated with 0.1% (w/v) Poly-L-Lysine (PLL, 70-150 kDa MW; Sigma-Aldrich) for 30 min at 25 °C. The coverslip was then washed twice with 2 ml of Milli-Q water and air-dried in the tissue culture hood for 2 h. Primary CD8^+^ T cells were fixed with 0.8% paraformaldehyde and 0.2% glutaraldehyde in PBS for 30 min at 4 °C. Next, 10^6^ T cells were added to a single well containing the PLL-coated glass slide. The plate was centrifuged for 1 min at 400 x *g* at 25 °C and incubated for a further 30 min at 4 °C. Next, the fixative was removed and replaced with 1 ml of PBS. Fixed T cells were kept at 4 °C until further processing. The PBS was removed, and the sample was washed 3 times with a 0.1 M phosphate buffer. Secondary fixation was performed in a 1% (w/v) OsO_4_ in 0.1 M phosphate buffer for 1 h at 4 °C. The sample was washed three times with Milli-Q water and dehydrated using step-wise dilutions of ethanol (50%, 70%, 90%, 95% and 100% (v/v)) incubating the sample for 5 min at 25 °C at each step. The 100% ethanol was replaced with 50% (v/v) hexamethyldisilazane (HMDS) in ethanol and then 100% (v/v) HMDS, and incubated for 2 min each time. HMDS was removed, and the sample was air-dried for 18 h. The sample was then sputter-coated with gold (10-15 nm) and imaged using a Zeiss Sigma 300 Scanning Electron Microscope with the SE2 detector in operation.

### Preparation of primary CD8^+^ T-cells and U-2 OS cells for live-cell imaging

For imaging of T cell/cancer cell interactions, 2.5×10^5^ U-2 OS cells in 400 μl of complete McCoy’s media were seeded in a glass-bottom 8-chamber coverslip (Ibidi, Cat# 80807). Optionally, the media was supplemented with 10 μM gp100 peptide (for ^Lo^U-2 OS_WT_ and ^Lo^U-2 OS_ICAM-1_). U-2 OS cells were cultured for 18 h. Prior to imaging, the media was removed, U-2 OS cells were washed and incubated with 0.2 μM Janelia Fluor 549 HaloTag Ligand (Promega, Cat# GA1110) for 15 min. Next, U-2 OS cells were washed twice and incubated in media for 10 min to remove the residual dye. Optionally, 10 nM N-Med ImmTAC or 10 nM Janelia Fluor 646-conjugated N-Med ImmTAC were included in the media. After the incubation, the U-2 OS cells were washed three times. Finally, 300 µl of media was added to U-2 OS cells. For imaging of calcium flux, the imaging media were replaced with PBS supplemented with 10 mM HEPES and 2 mM MgSO_4_.

Primary T cells were thawed the day before imaging and cultured in complete RPMI media. On the day of the experiment, dead T cells were removed by MACS using the dead removal kit (Miltenyi Biotec, Cat# 130090101), LS columns (Miltenyi Biotec, Cat# 130042401) and a manual magnetic separator (Miltenyi Biotec, Cat# 130091051) according to the manufacturer’s protocol. 1×10^6^ T cells were harvested and centrifuged at 200 x *g* for 2 min. The cells were washed and resuspended in 5 µg/ml silicon rhodamine- or AF488-conjugated anti-CD45 antibody and incubated for 5 min. Optionally, the imaging media was supplemented with 0.5% (v/v) Fluo4-AM (ThermoFisher, Cat# F14201) or 1 μg/ml CellMask Green Plasma Membrane Stain (ThermoFisher, C37608). Primary T cells expressing HaloTagged transgenes were labelled in 0.2 µM Janelia Fluor 646 HaloTag ligand for 15 min. Next, T cells were washed twice and resuspended in 50 μl of imaging media or PBS supplemented with 10 mM HEPES and 2 mM MgSO_4_ for calcium flux imaging.

The 8-chambered coverslip containing was placed on the microscope stage and 20-50 μl of T-cell suspension was gently added to the chamber containing the U-2 OS cells. The acquisition was initiated as soon as the first T cell became visible above the U-2 OS cell monolayer.

All media and buffers used for imaging were syringe-filtered (0.22 μm, PES) before use. All incubation and wash steps were performed using phenol red-free RPMI media supplemented 10 mM HEPES with unless specified otherwise. All incubation steps were carried out in a humidified incubator at 37 °C and 5% CO_2_. Imaging was performed at 37 °C unless stated otherwise. Imaging data was viewed and processed using ImageJ software.

### Epifluorescence widefield microscopy

The instrument was fitted with a 100x Plan Apo TIRF, NA 1.49 oil immersion objective (Nikon) mounted on an inverted microscopy body (Eclipse Ti2, Nikon) contained within an incubation chamber (DigitalPixel). The sample was illuminated using the following lasers: 488 nm (iBeam-SMART, Toptica), 561 nm (LaserBoxx, DPSS, Oxxius) and 641 nm (Obis Coherent). The laser beams were circularly polarised, collimated, expanded and aligned. The excitation light was directed through the centre of the back focal plane of the objective to generate widefield illumination. The emitted fluorescence passed through the objective lens, dichroic mirror (Di01-R405/488/561/635, Semrock), filters (FF01-520/44-25, BLP01-488R, LP02-568RS-25, FF01-587/35-25, FF01-692/40-25; Semrock) mounted in an automated filter wheel and a tube lens (1.5x). The signal was recorded by an electron-multiplying charge-coupled device (Evolve 512 Delta, Photometrics) with an electron multiplication gain of 250 ADU/photon operating in frame transfer mode. The effective pixel size was 107 nm. For imaging of 1G4-HaloTag CD8^+^ T-cells, the microscope was fitted with a Prime BSI Express sCMOS Camera (Teledyne Photometrics). The effective pixel size was 44 nm. Each image was acquired as a z-stack of 17 focal planes with an interplane interval of 200 nm. The three- or two-color images were acquired sequentially at a frequency of 0.05 Hz with 100 ms exposure time. The laser powers were optimised to reduce bleaching while maximising signal.

### Single-objective light sheet microscopy

Single-objective light sheet microscope based on the open-top eSPIM configuration as reported by Yang *et al*. was used^34^. The eSPIM instrument was fitted with a 60x water immersion objective (CFI Plan Apo IR 60XC WI, MRD07650, Nikon) mounted on an inverted microscope body (Eclipse Ti-U, Nikon) contained within an incubation chamber (DigitalPixel). The sample was illuminated with the following lasers: 561 nm (06-MLD-561-100, Cobolt) and 638 nm (06-MLD-638-180, Cobolt). The excitation light was filtered (MF559-34, Semrock; FF01-637/7-25, Thorlabs), attenuated, expanded and collimated. The laser lines were combined into a single excitation path. The excitation light was directed through a cylindrical lens to generate a Gaussian light sheet, and its optical properties were modulated by an adjustable slit. The Gaussian beam was then projected through the edge of the back focal plane of the primary objective to produce a tilted (30°) light-sheet illumination onto the sample. The emitted fluorescence was collected through the same primary objective and passed through a dichroic mirror (Di01-R406/488/561/635; Semrock), a secondary objective (CFI Plan Apo Lambda D 40× Air 0.95 NA, MRD70470, Nikon) and a tertiary glass-tipped objective (AMS-AGY v1.0, Calico). The emission light was directed into an optical splitter (OptoSplit III, Prior) to separate the fluorescence channels and projected onto distinct portions of a sCMOS camera chip (Prime 95B, Photometrics). The two-color images were acquired simultaneously with an exposure time of 30 ms at a frequency of 0.1 Hz. The effective pixel size was 120 nm. The images were deskewed and converted into focal stacks or 3D projections using ImageJ and a bespoke python code (github.com/Zui409/3D-Deskew-and-Reconstruction-for-single-objective-SPIM-data). For image deconvolution, Richardson-Lucy (RL) deconvolution with 10 iterations was performed on the deskewed images. Quantification of the number of T cells interacting with U-2 OS cell monolayers at t = 10 min was performed through visual inspection of the reconstructed images.

### Spinning-disk confocal microscopy

Olympus SoRa spinning disk was fitted with oil-immersion objectives (60x 1.5NA UPlan Apo and 100x 1.45NA UPlanApo) and mounted on an inverted microscope body (Olympus IX83). The sample was illuminated with the following lasers: 488 nm, 561 nm and 650 nm (Obis Coherent). The emitted fluorescence passed through the objective lens, dichroic mirror (D405/488/561/640; Chroma), filters (B525/50, B525/50, B685/40; Chroma). The out-of-focus light was removed using a 50 µm pinhole disk (Yokogawa) and the signal was recorded by the sCMOS camera chip (98B, Photometrics). Each image was acquired as a z-stack of 15-20 focal planes with an interplane interval of 250 nm. The pixel size was 110 nm. The imaging frequency varied depending on the size of the focal stack. Image acquisition was performed using Olympus CellSens Dimension 4.3.1 software.

3i intelligent imaging spinning disk microscope was fitted with a 100x Plan Apo TIRF, NA 1.46 oil immersion objective (Zeiss) mounted on an inverted microscope body (Zeiss Axio Observer 7). The sample was illuminated by the following lasers: 488 nm (LuxX), 561 nm (OBIS) and 638 nm (LuxX). The excitation light was field flattened (Yokogawa-Uniformizer) and relayed through a spinning disk unit (Yokogawa CSU-W1 T2, SoRa Dual Microlens Disk) and a dichroic mirror (FF01-440/521/607/700, Semrock). The emission light was filtered (FF01-525/45-25-STR, FF01-692/40-25-STR, FF02-617/ 73-25-STR; Semrock). The fluorescence signal was recorded using a sCMOS camera (Prime 89B, Teledyne Photometrics).

### Close-contact analysis

Quantitative analysis of close contacts was carried out using custom code based on code that was described previously (github.com/mkoerbel/contactanalysis_2D)^9^. Briefly, all close contacts in a field of view were segmented by the code via difference of gaussian filtering followed by hysteresis thresholding. The code then calculated a range of close contacts features. These features were then normalised according to the number of cells present in the field of view.

### CD45 exclusion analysis

Quantitative analysis of CD45 exclusion was carried out using ImageJ. Close contacts were segmented via difference of gaussian filtering, followed by thresholding via Otsu’s Method. The cell footprint was segmented via gaussian filtering followed by thresholding via Otsu’s Method. The segmented close contact mask and the segmented cell footprint mask were used to generate a third mask that segmented regions that were inside the cell footprint, but not inside the close contacts. The CD45 and Cell mask channels were normalised such that all pixel values fell between 1 and 0 to account for differences in fluorescence intensity between the two channels. The CD45 channel was then divided by the cell mask channel to account for possible artefactual exclusion/enrichment caused by cell membrane fluctuations. The normalised CD45/Cell mask image was then used to calculate exclusion using the formula 1 - average pixel intensity inside close contact (calculated using the close contact mask)/average pixel intensity outside close contacts (calculated using the third mask).

### Protein enrichment analysis

Quantitative analysis of protein enrichment was carried out using ImageJ. Close contacts were segmented via difference of gaussian filtering, followed by thresholding via Otsu’s Method. The cell footprint was segmented via gaussian filtering followed by thresholding via Otsu’s Method. The segmented close contact mask and the segmented cell footprint mask were used to generate a third mask that segmented regions that were inside the cell footprint, but not inside the close contacts. The close contact mask was used to calculate the average pixel intensity (in the channel of the protein of interest) inside the close contacts, while the third mask was used to calculate the average pixel intensity outside the close contacts. Protein enrichment was then calculated using the formula: average pixel intensity inside close contact/average pixel intensity outside close contacts.

### Stochastic simulations of close contacts at cell-cell interfaces

Spatial stochastic simulations to model TCR triggering were performed using Smoldyn 2.73^75^. The simulations were run using a bespoke, parallel Python script. Analysis and plotting were done using Python 3.10.17 with the packages NumPy^76^, pandas^77^, matplotlib^78^, and seaborn^79^. All scripts have been deposited on Github (github.com/mkoerbel/contactsimulations_TCRtriggering).

### General setup

A single close contact at a cell-cell interface was projected onto 2D with a close contact radius *r*_*CC*_ = 220nm in a circular simulation with boundary radius *r*_*sim*_ = 1 µm. We thereby assume that a) the only ‘inside’ of the close contact the opposing cell membrane are close enough for antigen to bind to the TCR, c) ‘outside’ the close contact extends along the cell membrane beyond *r*_*sim*_, modelled by imposing reflecting boundary conditions, d) all species diffuse freely both ‘inside’ and ‘outside, and, beside for CD45 (see Model 2), the close contact boundary does not impose a diffusion barrier, and e) close contacts can be considered in isolation for the purpose of describing TCR triggering. At time *t* = 0, all molecular species were randomly placed inside the simulation at their respective densities (Table S3). For CD45 in model 2, the numbers inside and outside the contact were adjusted to yield the desired exclusion value. The system was simulated for 30 s with timestep Δ*t* = 0.002 s.

Binding of antigen to the TCR was modelled as a pseudo-first order reaction (independent of the antigen diffusion speed):

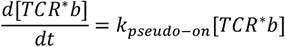

with *k*_*pseudo*–*on*_ = *k*_*on*_[*antigen*], [*species*] indicating the density of *ρ*_*species*_ in molecules/µm^2^ and *TCR*^*^ indicating any phosphorylation state. Unbinding followed first-order kinetics:

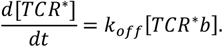

To reflect the experimental system, the *k*_*on*_ and *k*_*off*_ rates of the TCR-binding domain of N-Med ImmTAC were used (Table S1). Due to the high affinity of the pMHC-binding domain we considered this the rate determining binding site. The 3D rate constants were converted to 2D according to Huppa *et al*.^80^: 2*Dk*_*off*_ = 3*Dk*_*off*_, 2*DK*_*d*_ = 3*DK*_*d*_/1.239 ⋅ 10^−/^, and 2*Dk*_*on*_ = 2*Dk*_*off*_/2*DK*_*d*_.

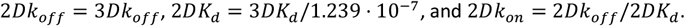

Phosphorylation of the multiple ITAMs on the TCR has previously been suggested to occur in random order^57^,we thus only track the total number of phosphorylations in both models.

All simulation parameters are summarised in Table S3.

### Setup Model 1

Model 1 is based on the dwell-time or kinetic segregation – kinetic proofreading (KS-KP) model^45,46^, which has been previously implemented in a similar way using Smoldyn to study the effect of TCR diffusion on triggering^17^. TCRs inside the close contact can be phosphorylated up to 9 times with rate *k*_*phos*_:

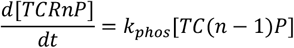

Upon diffusing out of the contact, the phosphorylations are immediately reset to *TCR*0*P*. A previous study using the kinetic proofreading model to study T-cell discrimination have estimated the number of proofreading steps to *n* = 3^81^. Experimental methods estimated the number of proofreading steps at the TCR level to *n* = 5^56^.

### Setup Model 2

Model 2 builds on the concepts in model 1 by explicitly modelling phosphorylation and dephosphorylation of the TCR assuming second-order kinetics:

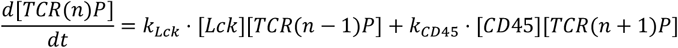

for 0<= *n* <= 5. These kinetics apply to both ImmTAC-bound and unbound TCRs. Because [*TCR*] < [*CD*45] and [*TCR*] < [*Lck*], and thus both enzymes do not operate at their *Vma*x (if described by Michaelis Menten kinetics), we argued that the enzyme reactions can be approximated by second order reactions to reduce the number of parameters in the model compared to Michaelis Menten kinetics. Kinases and phosphatases can react both inside and outside the close contact with TCR, allowing for background signaling outside the contact and re-entry of TCR into the contact without resetting its phosphorylation state. CD45 exclusion 𝜀 is modelled explicitly by setting the entry probability into the contact to 1 ™ 𝜀/100. To track the TCR phosphorylation states for analysis, TCR species were set to keep their serial number throughout reactions and saved together with the TCR state and coordinates every 10 timesteps.

### Notes for comparing Model 1 and Model 2

The phosphorylation rate *k*_*phos*_ in Model 1 comprises a net-phosphorylation rate, accounting for the kinetics of both Lck and CD45 inside the close contact. In terms of densities and rate constants used in Model 2, this scales as 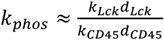. The CD45 density inside the contact *d*_*CD*45_ = *d*_*in*_ is required to define the exclusion value 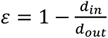 and consequently 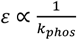. Therefore, when comparing Fig. 6 b nd f, the inverse relationship of the x-axes should be considered.

### Model Analysis

To consider a close contact as triggered, one TCR needs to reach a minimum number of *n*_*trig*_phosphorylations within the scan time. In model 1, because TCRs can only be dephosphorylated by leaving the close contact, the first time step any TCR in the simulation reaches n_trig_P is recorded as the triggering time *t*_*trig*_. The lifetime τ of a TCR phosphorylation state is thus capped by the TCR dwell-time inside the close contact. In model 2, because each TCR phosphorylation state is at equilibrium after a few time steps (Fig. S7F), a further requirement on the lifetime τ of the state was introduced, to model the necessity of a follow-up binding event. The lifetime of the TCR-5P phosphorylation state τ_5*P*_ was calculated for each TCR track in a simulation: A lifetime event started with the first time a TCR reached 5P, and ended when it got dephosphorylated. The triggering time *t*_*trig*_ was defined as the time point in a simulation when a single TCR reached 5P with τ_5*P*_ > τ_*min*_ for the first time. τ_*min*_ = 0.3 s was primarily used to reflect a possible triggering mechanism that relies on CD8 to be recruited: For a typical 3*Dk*_*on*_ = 10^5^ M_-1_s_-1_ of protein-protein association_82_, and a CD8 density of 300 µm_2 9_, after conversion to 2D rates as above, this results in a mean association time of 0.3 s. The mean phosphorylation time of Lck also is 0.3 s (*k*_*cat*_ = 3.41 s_-1_)_54_.

The lifetime events in model 2 were further categorised into ‘inside’ and ‘outside’ the close contact based on its location at the beginning of the event. Because the phosphorylation state of a TCR in model 2 is not reset upon diffusing across the close contact boundary, single lifetime events classified as ‘outside’ could show a dependence on the CD45 exclusion. In our analysis for triggering we take TCRs both inside and outside into account, and this becomes only relevant for plots that try to separate inside and outside TCR tracks.

We focused in model 2 on the 5P triggering endpoint, but note that the lifetimes will be longer if the triggering requirement is set to “at least 4P” in a simulation with maximum 5P (or generally more than 4 possible phosphorylation sites). This could increase the flexibility of the reaction system by having more phosphorylation sites on the TCR than are necessary for the next proofreading step.

