## Supplementary data for "T-cell signaling relies on partial CD45-exclusion at sub-micron sized cellular contacts"

### SUPPLEMENTARY FIGURES, MOVIE LEGENDS, AND TABLES

#### Supplementary figures

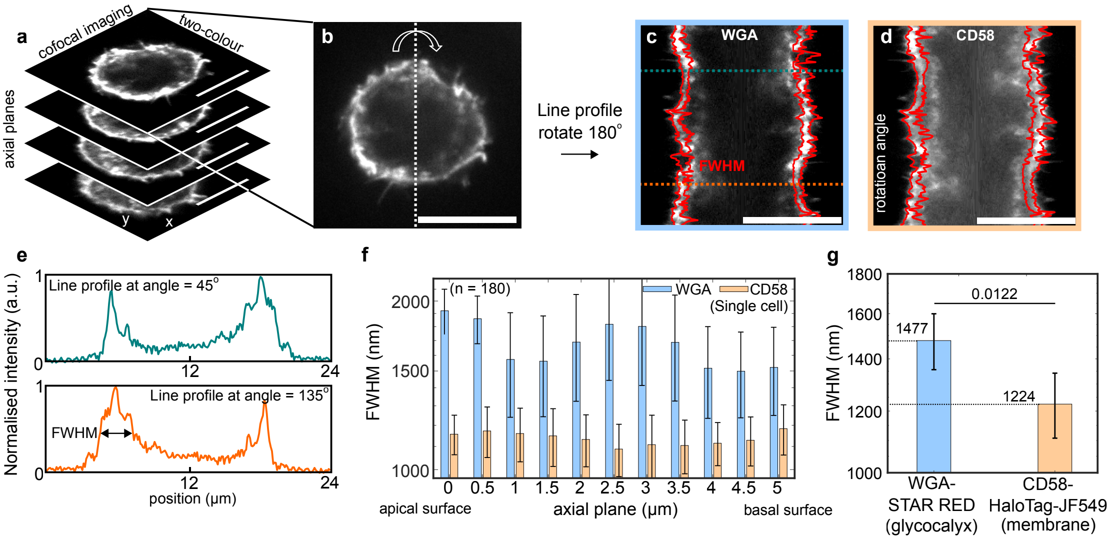


**Figure S1 | Quantification of glycocalyx thickness using custom radial line profile analysis.**

**a.** Representative 3D spinning-disk confocal image stack (single colour channel) of a U-2 OS cell line overexpressing CD58-HaloTag (referred to as U-2 $OS_{WT++}$). CD58-HaloTag was used as a membrane marker and was labelled using JF549 HaloTag ligand. Glycocalyx was labelled using STAR RED-conjugated Wheat Germ Agglutinin (WGA). **b.** Schematic of the analysis: in each 2D plane, a line profile is drawn through the cell's centre of mass and rotated in 1° increments over 180°, generating a continuous radial sampling of the entire 2D image plane. **c, d**. Example radial kymographs assembled from 180 radial line profiles for WGA (c) and CD58 (d). Red overlays indicate the computed full width at half maximum (FWHM) detected for each angle. **e.** Example intensity profiles at 45° and 135°, showing Gaussian fits used to extract σ and FWHM. **f.** Analysis across an entire single cell. This process was repeated for every axial plane, generating average FWHM values (mean ± SD; n = 180 profiles per plane) for WGA and CD58, measured across 11 axial planes spanning 5 µm from the apical to basal surface. **g.** Summary of FWHM values form multiple cells, averaged over entire 3D stacks across multiple cells (n = 6; total of 11,880 profiles). Data was compared using a two-tailed two-sample t test. All scale bars, 10 µm. All analyses were implemented in MATLAB. Error bars correspond to SD.

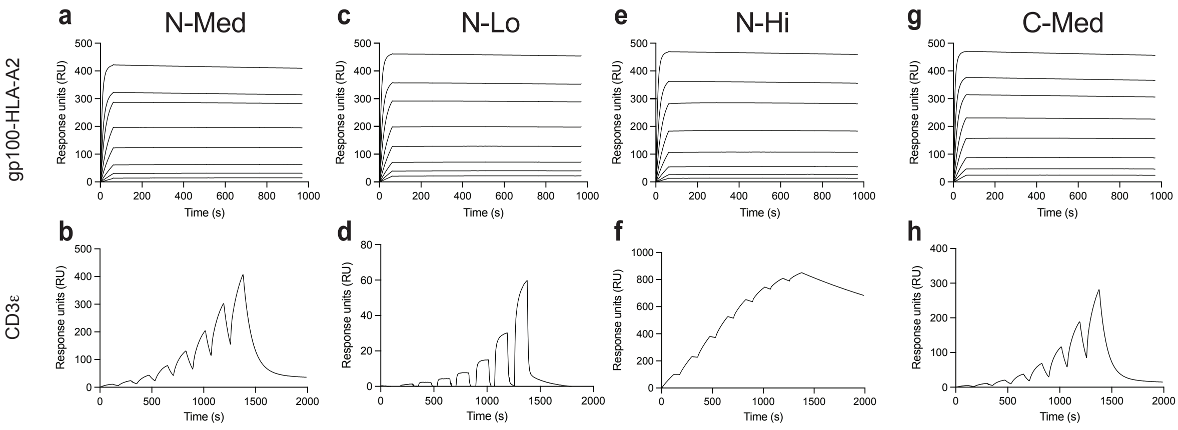


**Figure S2 | Surface plasmon resonance measurements of ImmTAC binding to their ligands. a, c, e, g** Multi-cycle SPR-based analysis of the ImmTACs binding to gp100-HLA-A2. 2xHis$_6$-tagged gp100-HLA-A2 was immobilised on an NTA chip and fluid-phase ImmTACs were injected as analytes. Twofold dilution series ranging from 1.56 nM to 200 nM was used. Sensorgrams were blank subtracted. All sensorgrams were fitted to a 1:1 Langmuir model to produce the kinetic parameters shown in Table S1. **b, d, f, h** Single-cycle SPR-based analysis of the ImmTACs binding to CD3ε. A biotinylated CD3εγ heterodimer was immobilised on a CAP chip and fluid-phase ImmTACs were injected as analytes. Twofold dilution series ranging from 6.25 nM to 800 nM was used. Sensorgrams were blank subtracted. All sensorgrams were fitted to a 1:1 Langmuir model to produce the kinetic parameters shown in Table S2.

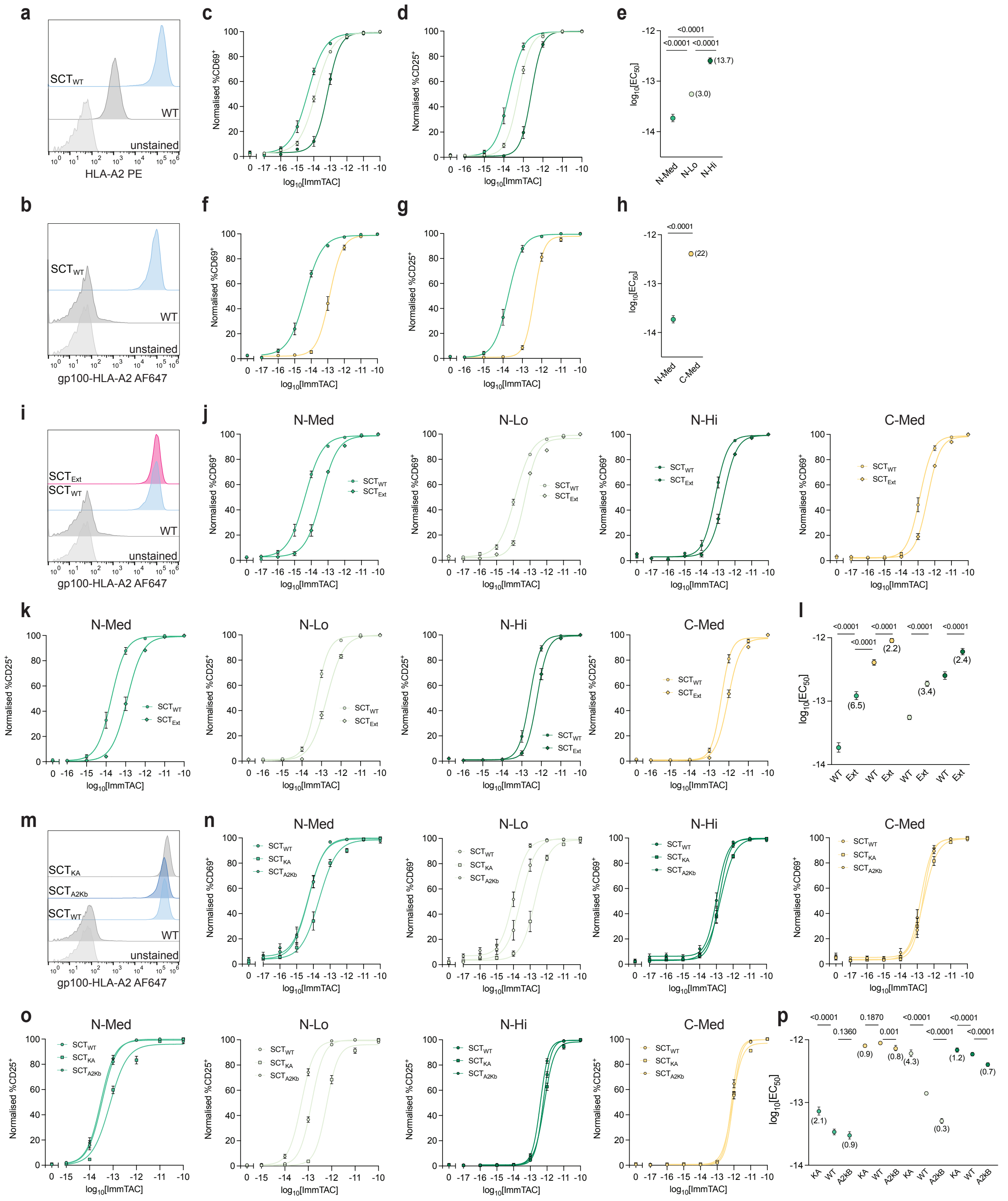


**Figure S3 | Creation of gp100-SCT U-2 OS cell lines and co-culture assays of CD8$^{+}$ T-cells with gp100-SCT U-2 OS cells showing pMHC-like properties of ImmTACs. a.** Flow-cytometry histogram showing expression of HLA-A2 in wild-type U-2 OS cells and U-2 OS cells expressing wild-type gp100-β$_2$M-HLA-A2 single chain trimer (SCT$_{WT}$). Staining was performed using a PE-conjugated anti-HLA-A2 antibody. **b.** Flow-cytometry histogram showing expression of gp100-HLA-A2 in wild-type U-2 OS cells and SCT$_{WT}$ U-2 OS cells. Staining was performed using an AF647-conjugated N-Med ImmTAC **c, d.** Dose-response curves showing the proportion of CD69$^{+}$ (c) and CD25$^{+}$ (d) by T cells following co-culture with SCT$_{WT}$-expressing

U-2 OS cells presenting N-terminal ImmTACs. Error bars correspond to SEM. **e.** $EC_{50}$ values measured after co-culturing $CD8^+$ T-cells with $SCT_{WT}$-expressing U-2 OS cells presenting N-terminal ImmTACs. Values in brackets indicate the fold shift in $EC_{50}$ values relative to N-Med. **f, g.** Dose-response curves showing the proportion of $CD69^+$ (f) and $CD25^+$ (g) T cells following co-culture with $SCT_{WT}$-expressing U-2 OS cells presenting N-Med and C-Med ImmTACs. Error bars correspond to SEM. **h.** $EC_{50}$ values measured after co-culturing $CD8^+$ T-cells with $SCT_{WT}$ U-2 OS cells presenting N-Med and C-Med ImmTACs. Values in brackets indicate the fold shift in $EC_{50}$ values relative to N-Med. **i.** Flow-cytometry histogram showing the matched expression of gp100-HLA-A2 in U-2 OS cells expressing $SCT_{WT}$ and $SCT_{EXT}$. Staining was performed using an AF647-conjugated N-Med ImmTAC. **j, k.** Dose-response curves showing the proportion of $CD69^+$ (j) and $CD25^+$ (k) T cells following co-culture with $SCT_{WT}$ and $SCT_{EXT}$ U-2 OS cells presenting all ImmTACs. Error bars correspond to SEM. **l.** $EC_{50}$ values measured after co-culturing $CD8^+$ T-cells with $SCT_{WT}$ and $SCT_{EXT}$ U-2 OS cells presenting all ImmTACs. Values in brackets indicate the fold shift in $EC_{50}$ values relative to $SCT_{WT}$. **m.** Flow-cytometry histogram showing the matched expression of gp100-HLA-A2 in U-2 OS cells expressing $SCT_{WT}$, $SCT_{KA}$, and $SCT_{A2kB}$. Staining was performed using an AF647-conjugated N-Med ImmTAC **n, o.** Dose-response curves showing the proportion of $CD69^+$ (n) and $CD25^+$ (o) T cells following co-culture with $SCT_{WT}$, $SCT_{KA}$, and $SCT_{A2kB}$ U-2 OS cells presenting all ImmTACs. Error bars correspond to SEM. **p**. $EC_{50}$ values measured after co-culturing $CD8^+$ T-cells with $SCT_{WT}$, $SCT_{KA}$, and $SCT_{A2kB}$ U-2 OS cells presenting all ImmTACs. Values in brackets indicate the fold shift in $EC_{50}$ values relative to $SCT_{WT}$. All $EC_{50}$ values were determined by fitting a Hill function to dose-response curves showing the proportion of $CD25^+$ T cells. $EC_{50}$ values were compared using an F-test of the normalised data. All error bars for the $EC_{50}$ values correspond to 95% CI. All co-culture assays were performed as three biological repeats in triplicate. Data were collected using $CD8^+$ T-cells isolated from at least two donors.

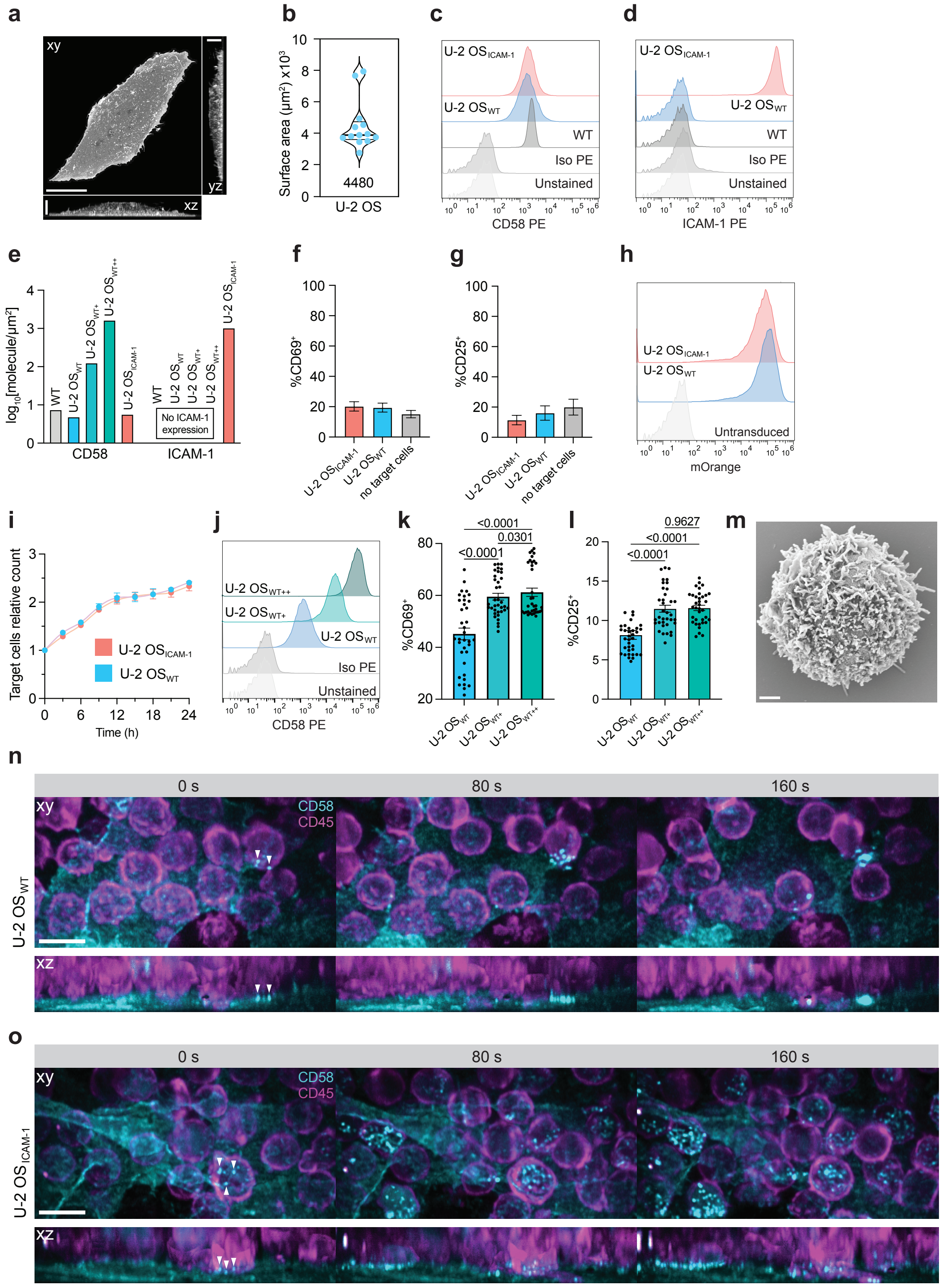


**Figure S4 | Creation and characterisation of U-2 OS cell lines for imaging of close contacts, and epi-illumination selective-plane illumination microscopy of T cell/U-2 OS cell interactions. a.** Point-scanning confocal image of a fixed U-2 OS cell stained with a membrane dye. Orthogonal planes (xy, xz, and yz) are shown as maximal-intensity projections of the focal stacks. Scale bars, 20 µm (xy) and 5 µm (xz and yz). **b.**

Estimated surface area of the U-2 OS cell surface calculated from images such as those shown in (a); n = 13. **c.** Flow-cytometry histogram showing matched expression of CD58 in wild-type U-2 OS, U-2 $OS_{WT}$, and U-2 $OS_{ICAM-1}$ cells. Staining was performed using a PE-conjugated anti-CD58 antibody **d.** Flow-cytometry histogram showing expression of ICAM-1 in wild-type U-2 OS, U-2 $OS_{WT}$, and U-2 $OS_{ICAM-1}$ cells. Staining was performed using a PE-conjugated anti-ICAM-1 antibody **e.** Density of CD58 and ICAM-1 on the surface of wild-type U-2 OS, U-2 $OS_{WT}$, U-2 $OS_{WT+}$, U-2 $OS_{WT++}$ and U-2 $OS_{ICAM-1}$ cells calculated using data from (a-d and j). **f, g.** Proportion of $CD69^+$ (g) and $CD25^+$ (h) T-cells following co-culture with U-2 $OS_{WT}$ and U-2 $OS_{ICAM-1}$ cells or without target cells. n = 3; error bars correspond to SEM. Co-culture assays were performed as three biological repeats in triplicate. Data were collected using $CD8^+$ T-cells isolated from at least two donors. **h**. Flow-cytometry histograms showing expression of mOrange by U-2 $OS_{WT}$ and U-2 $OS_{ICAM-1}$ cells for killing assays. **i**. Killing of U-2 OS cells by $CD8^+$ T-cells over time, measured using Incucyte imaging. Killing assays were performed using $CD8^+$ T-cells isolated from three donors in triplicate. Error bars correspond to SEM. **j.** Flow-cytometry histogram showing expression of CD58 in U-2 $OS_{WT}$, U-2 $OS_{WT+}$ and U-2 $OS_{WT++}$ cells. Staining was performed using a PE-conjugated anti-CD58 antibody. **k, l.** Proportion of $CD69^+$ (k) and $CD25^+$ (l) T-cells following co-culture with U-2 $OS_{WT}$, U-2 $OS_{WT+}$ and U-2 $OS_{WT++}$ cells in the absence of TCR ligand. n = 36; error bars correspond to SEM. Co-culture assays were performed as three biological repeats. Data were collected using $CD8^+$ T-cells isolated from at least two donors. Data were compared using one-way ANOVA with Geisser-Greenhouse correction. **m.** Scanning electron microscopy image of a $CD8^+$ T-cell. Scale bar, 0.5 µm. **n, o.** Epi-illumination selective-plane illumination imaging of $CD8^+$ T-cells interacting with U-2 $OS_{WT}$ (n) and U-2 $OS_{ICAM-1}$ (o) cells. Orthogonal planes (xy and xz) are shown as maximal-intensity projections. Scale bar, 10 µm.

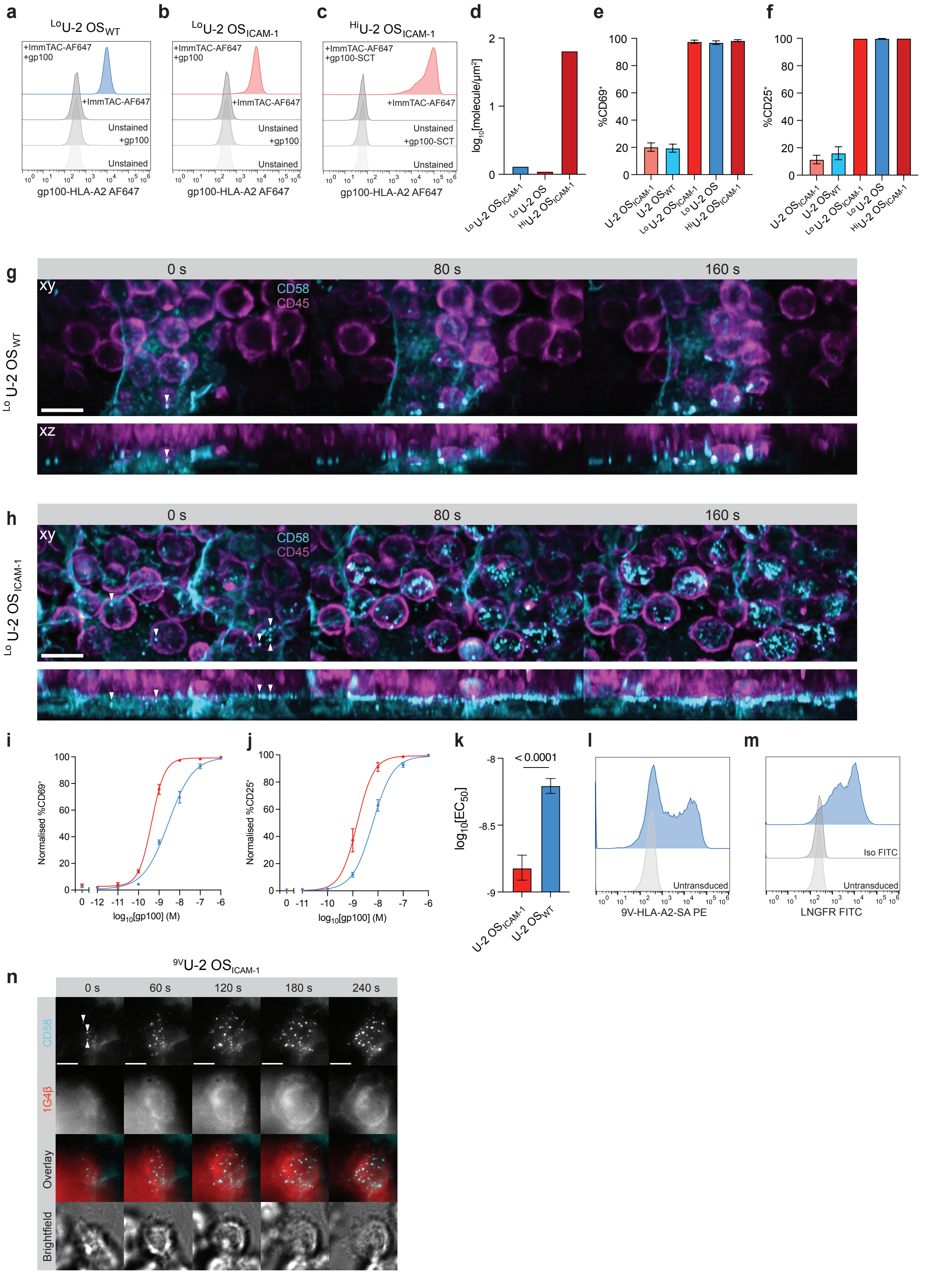


**Figure S5 | Characterisation of ImmTAC densities on the U-2 OS cell surface, epi-illumination selective-plane illumination microscopy of T cell/U-2 OS cell interactions, and effect of ICAM-1 expression on T-cell activation. a-c.** Flow-cytometry histograms showing N-Med ImmTAC levels on the surface gp100-pulsed $^{Lo}$U-2 OS$_{WT}$ (a), $^{Lo}$U-2 OS$_{ICAM\text{-}1}$ (b) cells and SCT$_{WT}$-expressing $^{Hi}$U-2 OS$_{ICAM\text{-}1}$ cells (c). **d**. Densities of

N-Med ImmTAC on the surface of $^{Lo}$U-2 $OS_{WT}$, $^{Lo}$U-2 $OS_{ICAM-1}$, and $^{Hi}$U-2 $OS_{ICAM-1}$ cells calculated using data from (a-c and Fig. S4a,b) **e, f**. Proportion of $CD69^+$ (f) and $CD25^+$ (g) T cells following co-culture with with U-2 $OS_{WT}$, U-2 $OS_{ICAM-1}$, $^{Lo}$U-2 $OS_{WT}$, $^{Lo}$U-2 $OS_{ICAM-1}$, and $^{Hi}$U-2 $OS_{ICAM-1}$ cells. n = 3; error bars correspond to SEM. Co-culture assays were performed as three biological repeats in triplicate. Data was collected using $CD8^+$ T-cells isolated from at least two donors. **g, h.** Epi-illumination selective-plane illumination imaging of $CD8^+$ T-cells interacting with $^{Lo}$U-2 $OS_{WT}$ (g) and $^{Lo}$U-2 $OS_{ICAM-1}$ (h) cells presenting N-Med ImmTAC. Orthogonal planes (xy and xz) are shown as maximal-intensity projections. Scale bar, 10 µm. **i, j.** Dose-response curves showing the fractions of $CD69^+$ (i) and $CD25^+$ (j) T-cells following co-culture with U-2 $OS_{WT}$ and U-2 $OS_{ICAM-1}$ presenting N-Med ImmTAC. Error bars correspond to SEM. **k.** $EC_{50}$ values measured after co-culturing $CD8^+$ T-cells with U-2 $OS_{WT}$ and U-2 $OS_{ICAM-1}$ cells presenting N-Med ImmTAC. $EC_{50}$ values were determined by fitting a Hill function to dose-response curves showing the proportion of $CD25^+$ T-cells. $EC_{50}$ values were compared using an F-test of the normalised data. Error bars for the $EC_{50}$ values correspond to 95% CI. Co-culture assays were performed as three biological repeats in triplicate. Data was collected using $CD8^+$ T-cells isolated from at least two donors. **l**. Flow-cytometry histograms showing expression of 1G4 TCR in primary $CD8^+$ T-cells. Staining was performed using a PE-conjugated 9V-HLA-A2-streptavidin (SA) tetramer **m**. Flow-cytometry histograms showing expression of LNGFR in primary 1G4 $CD8^+$ T-cells. LNGFR is contained within the polycistronic 1G4αβ construct and thus acts as an expression reporter. Staining was performed using a FITC-conjugated anti-LNGFR antibody. **n.** Epifluorescence widefield imaging of 1G4-HaloTag $CD8^+$ T-cells interacting with a monolayer of $^{9V}$U-2 $OS_{ICAM-1}$ presenting 9V-HLA-A2 antigen. White arrowheads indicate the first detectable close-contacts marked by accumulation of fluorescent CD58-HaloTag expressed by the U-2 OS cells. Images are displayed as maximum-intensity projections. Scale bars, 5 µm.

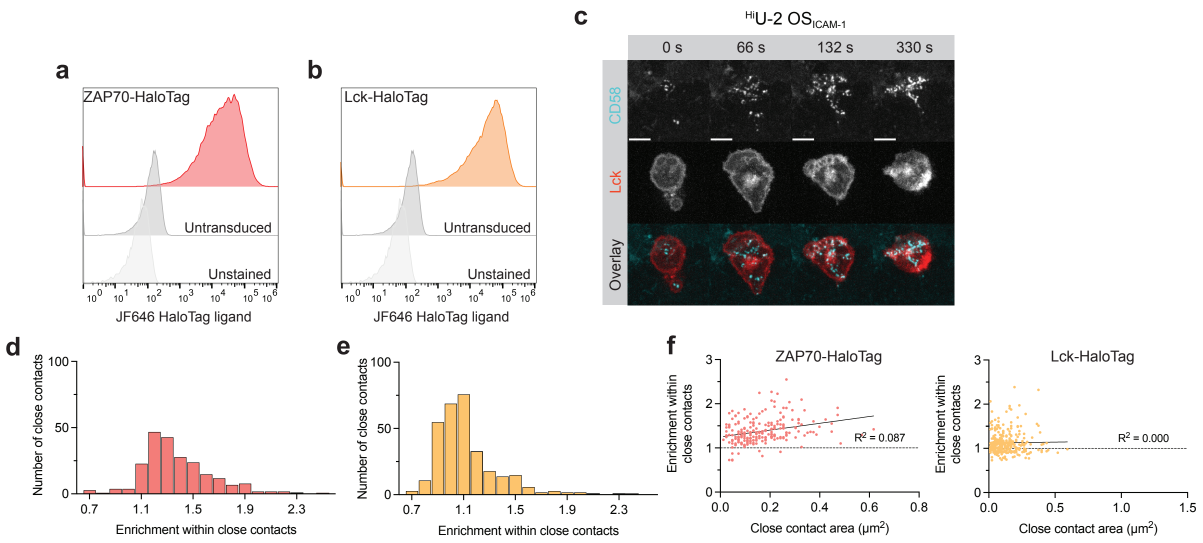


**Figure S6 | Creation of Lck-HaloTag and ZAP70-HaloTag CD8⁺ T-cells and analysis of their interactions with U-2 OS cells. a, b.** Flow-cytometry histogram showing expression of ZAP70-HaloTag (a) and Lck-HaloTag (b) in CD8⁺ T-cells. **c.** Spinning disk confocal imaging of interactions of Lck-HaloTag CD8⁺ T-cells with a monolayer of $^{Hi}$U-2 OS$_{ICAM-1}$ cells presenting N-Med ImmTAC. Scale bars, 5 µm. **d, e**. Histograms showing the enrichment levels for ZAP70-HaloTag (d) and Lck-HaloTag (e) at close contacts at the interface of CD8⁺ T-cells and $^{Hi}$U-2 OS$_{ICAM-1}$ cells presenting N-Med ImmTACs. **f.** Plots showing enrichment levels of ZAP70-HaloTag and Lck-HaloTag versus close-contact size.

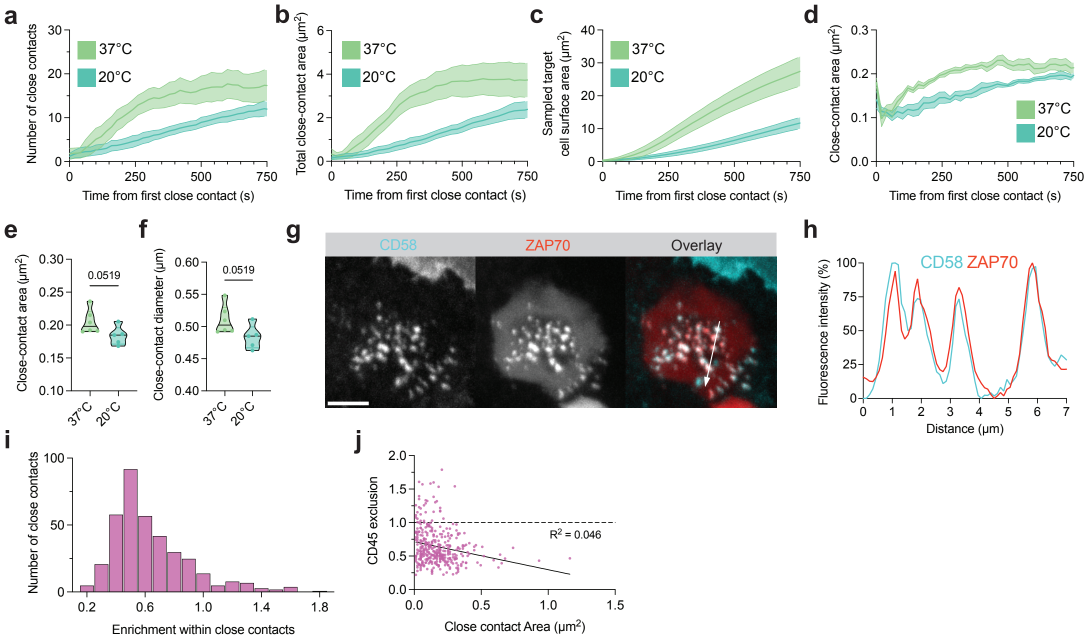


**Figure S7 | Interactions of CD8$^+$ T-cells with U-2 OS cells at reduced temperature. a-f.** Image-based analysis of CD8$^+$ T-cells interacting with $^{Lo}$U-2 OS$_{ICAM-1}$ cells presenting N-Med ImmTAC at 20° C and 37° C. Analysis was performed using epifluorescence widefield imaging data **a.** Average number of close contacts formed per T cell over time. **b.** Average summed area of close contacts per T cell over time. **c.** Average cumulative area of the target-cell surface scanned by a single T-cell over time. **d.** Average area of individual close contacts area versus time. **e, f.** Average area (e) and diameter (f) of all close contacts. **g.** Spinning-disk confocal imaging of interactions of ZAP70-HaloTag-expressing CD8$^+$ T-cells with a monolayer of $^{Hi}$U-2 OS$_{ICAM-1}$ cells presenting N-Med ImmTAC at 20° C. Scale bar, 5 µm. **h**. The min/max normalised intensity line profile was taken along the direction of the white arrow in (g). **i**. Histogram showing the CD45 exclusion levels at close contacts for CD8$^+$ T-cells interacting with $^{Lo}$U-2 OS$_{ICAM-1}$ cells presenting N-Med ImmTAC. **j.** Plot of CD45 exclusion levels versus close-contact size. Analysis in (i) and (j) was performed using the spinning-disk microscopy data collected at 20 °C.

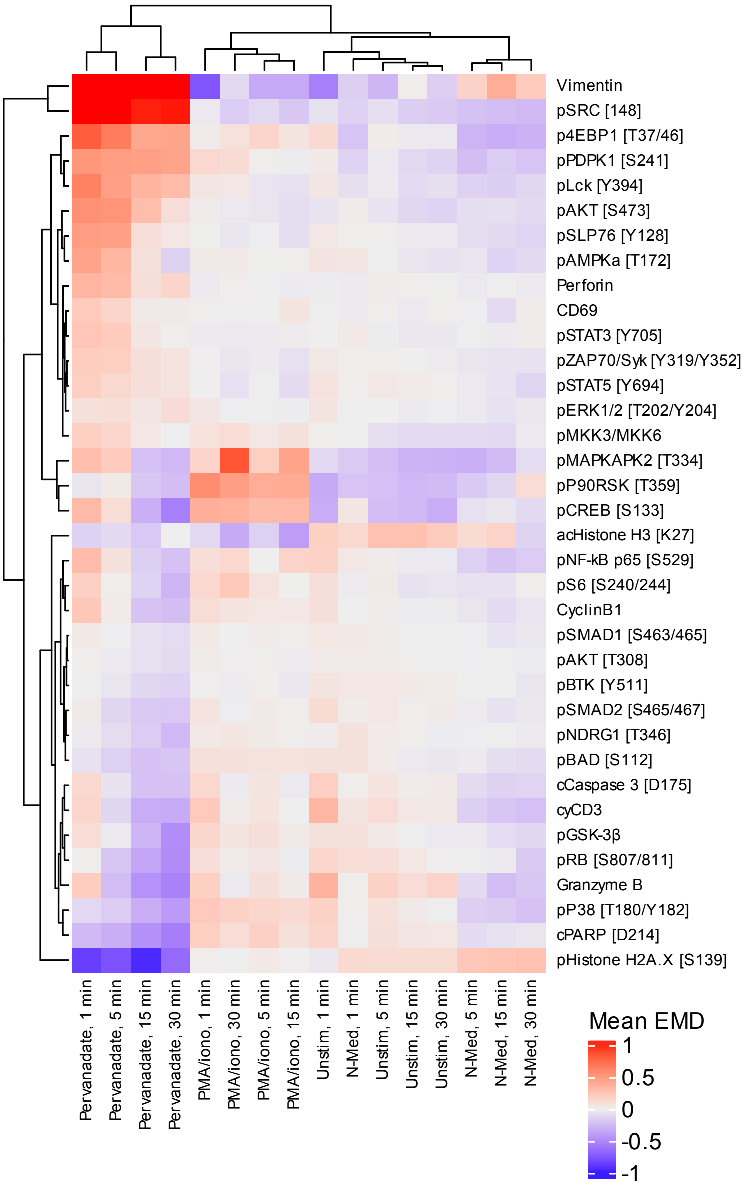


**Figure S8** | **Extended analysis of the CyTOF dataset**. Heatmap showing the mean of Earth Mover's Distance (EMD) scores for each marker across different timepoints. T cells from two donors were left unstimulated or stimulated with ImmTac, PMA/ionomycin, or pervanadate. Fluorescence intensity was measured by CyTOF at 1, 5, 15, and 30 minutes post-stimulation using a panel of 44 markers. Eight control markers were excluded from downstream analysis. Raw data were preprocessed using FlowJo, and analyzed with the CyGNAL package using default settings. To measure similarity between different treatment/timepoint conditions, EMD scores were calculated and normalized across the entire dataset. Four replicates for each condition. Hierarchical clustering was performed using Euclidean distance and complete linkage.

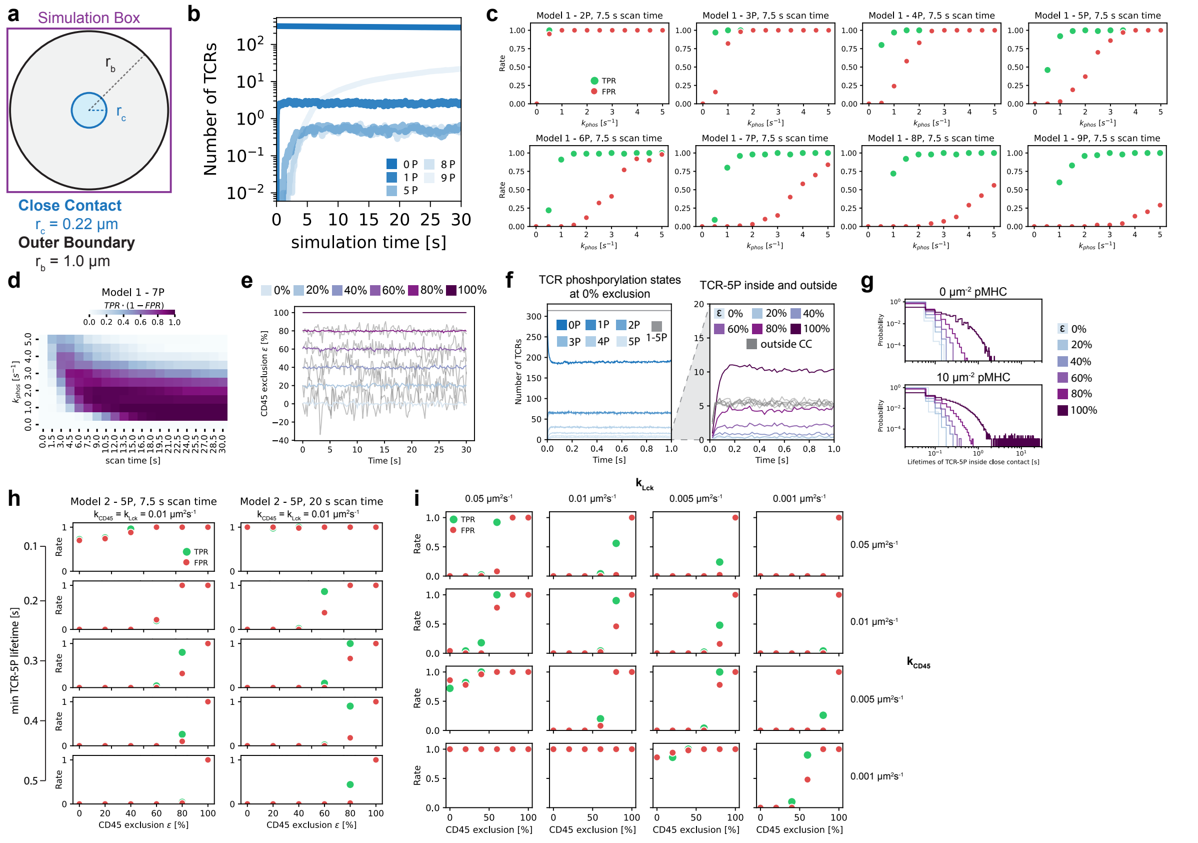


**Figure S9 | Extended plots and parameter spaces for stochastic simulations of model 1 and 2. a.** Schematic of the simulation box and close-contact size of Model 1 and 2. **b-d.** Supplementary data obtained with model 1. **b.** TCR dynamics versus time for $k_{phos}$ = 2. The numbers for bound and unbound TCR were summed, and the average of 100 simulations is shown. Only some of the 9 phosphorylation states are shown for better visibility. **c.** True positive rate (TPR) and false positive rate (FPR) for varying $k_{phos}$ and P, for 7.5 s scan time, calculated from 100 simulations. **d.** Heatmap indicating the TPR*(1-FPR) value for 7 phosphorylations varying $k_{phos}$ and scan time. In comparison to Fig. 6d, a wider range of $k_{phos}$ values for a given scan time yield a higher summary statistic. **e-h.** Supplementary results for Model 2 based on 50 simulations with $k_{CD45}$ = $k_{Lck}$ = 0.01 μm$^2$s$^{-1}$. **e.** Average measured CD45 exclusion for simulations with varying CD45 exclusion setpoint and 0 μm$^{-2}$ pMHC. The grey lines represent the exclusion over time for a single simulation, highlighting the stochasticity of the number of CD45 molecules inside the close contact. **f.** Average number of TCR in each phosphorylation state over time. Left: for $\varepsilon = 0$ the number of TCRs with 1-5P over the first second of the simulation are shown. The numbers remain constant until the end of the simulation. TCRs inside and outside the close contact have been pooled. Right: number of TCR-5P inside (colored) and outside (grey) the close contact over time for changing exclusion values. The higher the exclusion, the higher the number of TCR-5P inside the close contact. TCR-5P numbers outside the close contact are independent of exclusion. At a density of 100 μm$^{-2}$, there are 15 TCRs on average inside the close contact. **g.** Lifetime distributions of the TCR-5P state inside the close contact for different exclusion values and pMHC densities of 0 $\mu m^{-2}$ (top) or 10 $\mu m^{-2}$ (bottom). The lifetimes of individual TCR-5Ps of 50 simulations have been aggregated. **h.** TPR and FPR for 7.5 and 20 s scan time, while varying the minimum lifetime requirement of TCR-5P for triggering. For a 7.5 s scan time a lifetime of 0.3 s results in the highest separation of TPR and FPR at 80% exclusion. Increasing the scan time to 20 s, broadens the optimal lifetimes to 0.2-0.4 s for 60-80% exclusion. **i.** $k_{CD45}$ and $k_{Lck}$ parameter space for Model 2 with a lifetime cutoff of 0.3 s and 10 s scan time, showing the TPR and FPR. These are exemplary for the underlying data shown in Fig. 6h.

**Supplementary movie legends**

**Supplementary Movie 1 | Interactions of primary CD8$^+$ T cells with U-2 OS$_{WT}$ cells.** Epifluorescence widefield microscopy movie of a CD8$^+$ T cell interacting with a monolayer of U-2 OS$_{WT}$ cells. T cells were labelled with a SiR-conjugated anti-CD45 antibody acting as a cell membrane proxy. Close contacts are marked by the puncta of CD58-HaloTag accumulation. CD58-HaloTag was expressed by U-2 OS$_{WT}$ cells and labelled using JF549 HaloTag ligand. Movie is a maximal-intensity projection of the focal stack. Scale bar, 5 µm.

**Supplementary Movie 2 | Interactions of primary CD8$^+$ T cells with U-2 OS$_{ICAM-1}$ cells.** Epifluorescence widefield microscopy movie of a CD8$^+$ T cell interacting with a monolayer of U-2 OS$_{ICAM-1}$ cells. In the presence of ICAM-1, T cells form numerous, well-separated close contacts with target cells. T cells were labelled with a SiR-conjugated anti-CD45 antibody acting as a cell membrane proxy. Close contacts are marked by the puncta of CD58-HaloTag accumulation. CD58-HaloTag was expressed by U-2 OS$_{ICAM-1}$ cells and labelled using JF549 HaloTag ligand. Movie is a maximal-intensity projection of the focal stack. Scale bar, 5 µm.

**Supplementary Movie 3 | Overview of interactions between primary CD8$^+$ T cells and U-2 OS$_{WT}$ cells.** Epi-illumination selective-plane illumination microscopy movie of CD8$^+$ T cells interacting with a monolayer of U-2 OS$_{WT}$ cells. T cells were labelled with a SiR-conjugated anti-CD45 antibody acting as a cell membrane proxy. Close contacts are marked by the puncta of CD58-HaloTag accumulation. CD58-HaloTag was expressed by U-2 OS$_{WT}$ cells and labelled using JF549 HaloTag ligand. Orthogonal planes (xy and xz) are shown as maximal-intensity projections. Scale bar, 10 µm

**Supplementary Movie 4 | Overview of interactions between primary CD8$^+$ T cells and U-2 OS$_{ICAM-1}$ cells.** Epi-illumination selective-plane illumination microscopy movie of CD8$^+$ T cells interacting with a monolayer of U-2 OS$_{ICAM-1}$ cells. T cells were labelled with a SiR-conjugated anti-CD45 antibody acting as a cell membrane proxy. Close contacts are marked by the puncta of CD58-HaloTag accumulation. CD58-HaloTag was expressed by U-2 OS$_{ICAM-1}$ cells and labelled using JF549 HaloTag ligand. Orthogonal planes (xy and xz) are shown as maximal-intensity projections. Scale bar, 10 µm

**Supplementary Movie 5 | Interactions of primary CD8$^+$ T cells with $^{Lo}$U-2 OS$_{WT}$ cells.** Epifluorescence widefield microscopy movie of a CD8$^+$ T cell interacting with a monolayer of ImmTAC-presenting $^{Lo}$U-2 OS$_{WT}$ cells. T cells were labelled with a SiR-conjugated anti-CD45 antibody acting as a cell membrane proxy. Close contacts are marked by the puncta of CD58-HaloTag accumulation. CD58-HaloTag was expressed by $^{Lo}$U-2 OS$_{WT}$ cells and labelled using JF549 HaloTag ligand. Movie is a maximal-intensity projection of the focal stack. Scale bar, 5 µm.

**Supplementary Movie 6 | Interactions of primary CD8$^+$ T cells with $^{Lo}$U-2 OS$_{ICAM-1}$ cells.** Epifluorescence widefield microscopy movie of a CD8$^+$ T cell interacting with a monolayer of ImmTAC-presenting $^{Lo}$U-2 OS$_{ICAM-1}$ cells. T cells were labelled with a SiR-conjugated anti-CD45 antibody acting as a cell membrane proxy. Close contacts are marked by the puncta of CD58-HaloTag accumulation. CD58-HaloTag was expressed by $^{Lo}$U-2 OS$_{ICAM-1}$ cells and labelled using JF549 HaloTag ligand. Movie is a maximal-intensity projection of the focal stack. Scale bar, 5 µm.

**Supplementary Movie 7 | Overview of interactions between primary CD8$^+$ T cells and $^{Lo}$U-2 OS$_{WT}$ cells.** Epi-illumination selective-plane illumination microscopy movie of CD8$^+$ T cells interacting with a monolayer of ImmTAC-presenting $^{Lo}$U-2 OS$_{WT}$ cells. T cells were labelled with a SiR-conjugated anti-CD45 antibody acting as a cell membrane proxy. Close contacts are marked by the puncta of CD58-HaloTag accumulation. CD58-HaloTag was expressed by $^{Lo}$U-2 OS$_{WT}$ cells and labelled using JF549 HaloTag ligand. Orthogonal planes (xy and xz) are shown as maximal-intensity projections. Scale bar, 10 µm

**Supplementary Movie 8 | Overview of interactions between primary CD8$^+$ T cells and $^{Lo}$U-2 OS$_{ICAM-1}$ cells.** Epi-illumination selective-plane illumination microscopy movie of CD8$^+$ T cells interacting with a monolayer of ImmTAC-presenting $^{Lo}$U-2 OS$_{ICAM-1}$ cells. T cells were labelled with a SiR-conjugated anti-CD45 antibody acting as a cell membrane proxy. Close contacts are marked by the puncta of CD58-HaloTag accumulation. CD58-HaloTag was expressed by $^{Lo}$U-2 OS$_{ICAM-1}$ cells and labelled using JF549 HaloTag ligand. Orthogonal planes (xy and xz) are shown as maximal-intensity projections. Scale bar, 10 µm

**Supplementary Movie 9 | Interactions of primary 1G4 CD8$^+$ T cells with $^{9V}$U-2 OS$_{ICAM-1}$ cells.** Epifluorescence widefield microscopy movie of 1G4-HaloTag CD8$^+$ T-cells interacting with a monolayer of $^{9V}$U-2 OS$_{ICAM-1}$ cells presenting 9V-HLA-A2. 1G4β-HaloTag fusion protein was labelled using JF646-HaloTag ligand. Close contacts are marked by the puncta of CD58-HaloTag accumulation. CD58-HaloTag was expressed by $^{9V}$U-2 OS$_{ICAM-1}$ cells and labelled using JF549 HaloTag ligand. Movie is a maximal-intensity projection of the focal stack. Scale bar, 5 µm.

**Supplementary Movie 10 | ImmTAC accumulates within the close contacts.** Spinning-disk confocal microscopy movie of a CD8$^+$ T cell interacting with a monolayer of $^{Hi}$U-2 OS$_{ICAM-1}$ cells presenting a JF646-conjugated ImmTAC. Close contacts are marked by the puncta of CD58-HaloTag accumulation. CD58-HaloTag was expressed by $^{Hi}$U-2 OS$_{ICAM-1}$ cells and labelled using JF549 HaloTag ligand. Movie is a maximal-intensity projection of the focal stack. Scale bar, 5 µm.

**Supplementary Movie 11 | ZAP70 is recruited to the close contacts.** Spinning-disk confocal microscopy movie of a CD8$^+$ T cell expressing ZAP70-HaloTag fusion protein interacting with a monolayer of ImmTAC-presenting $^{Hi}$U-2 OS$_{ICAM-1}$ cells. ZAP70-HaloTag fusion protein was labelled using JF646-HaloTag ligand. Close contacts are marked by the puncta of CD58-HaloTag accumulation. CD58-HaloTag was expressed by $^{Hi}$U-2 OS$_{ICAM-1}$ cells and labelled using JF549 HaloTag ligand. Movie is a maximal-intensity projection of the focal stack. Scale bar, 5 µm.

**Supplementary Movie 12 | Lck is uniformly distributed across the T-cell membrane during close-contact formation.** Spinning-disk confocal microscopy movie of a CD8$^+$ T cell expressing Lck-HaloTag fusion protein interacting with a monolayer of ImmTAC-presenting $^{Hi}$U-2 OS$_{ICAM-1}$ cells. Lck-HaloTag fusion protein was labelled using JF646-HaloTag ligand. Close contacts are marked by the puncta of CD58-HaloTag accumulation. CD58-HaloTag was expressed by $^{Hi}$U-2 OS$_{ICAM-1}$ cells and labelled using JF549 HaloTag ligand. Movie is a maximal-intensity projection of the focal stack. Scale bar, 5 µm.

**Supplementary Movie 13 | CD8$^+$ T cells form close contact with $^{Lo}$U-2 OS$_{WT}$ cells before intracellular calcium flux.** Epifluorescence widefield microscopy movie of a CD8$^+$ T cell interacting with a monolayer of $^{Lo}$U-2 OS$_{WT}$ cells. Single close contact is visible before intracellular calcium flux. T cells were labelled with Fluo4 AM and SiR-conjugated anti-CD45 antibody acting as a cell membrane proxy. Close contacts are marked by the puncta of CD58-HaloTag accumulation. CD58-HaloTag was expressed by $^{Lo}$U-2 OS$_{WT}$ cells and labelled using JF549 HaloTag ligand. Movie is a maximal-intensity projection of the focal stack. Scale bar, 5 µm.

**Supplementary Movie 14 | CD8$^+$ T cells form close contact with $^{Lo}$U-2 OS$_{ICAM-1}$ cells before intracellular calcium flux.** Epifluorescence widefield microscopy movie of CD8$^+$ T-cells interacting with a monolayer of $^{Lo}$U-2 OS$_{ICAM-1}$ cells. Single close contact is visible before intracellular calcium flux. T cells were labelled with Fluo4 AM and a SiR-conjugated anti-CD45 antibody acting as a cell membrane proxy. Close contacts are marked by the puncta of CD58-HaloTag accumulation. CD58-HaloTag was expressed by $^{Lo}$U-2 OS$_{WT}$ cells and labelled using JF549 HaloTag ligand. Movie is a maximal-intensity projection of the focal stack. Scale bar, 5 µm.

**Supplementary tables**

**Table S1:** Kinetic parameters for ImmTAC binding to gp100-HLA-A2 ligand

| **ImmTAC** | **Replicate** | **$k_{on}$ ($M^{-1}s^{-1}$)** | **$K_{off}$ ($s^{-1}$)** | **$K_D$ (pM)** |
|---|---|---|---|---|
| | 1 | $4.45\times10^{5}$ | $3.73\times10^{-5}$ | 83.8 |
| N-Med | 2 | $1.31\times10^{6}$ | $3.52\times10^{-5}$ | 26.8 |
| | Average | $8.78\times10^{5}$ | $3.63\times10^{-5}$ | 55.3 |
| | 1 | $3.85\times10^{5}$ | $3.20\times10^{-5}$ | 83.0 |
| N-Lo | 2 | $9.64\times10^{5}$ | $2.93\times10^{-5}$ | 30.4 |
| | Average | $6.75\times10^{5}$ | $3.07\times10^{-5}$ | 56.7 |
| | 1 | $3.20\times10^{5}$ | $3.77\times10^{-5}$ | 118 |
| N-Hi | 2 | $1.45\times10^{6}$ | $3.56\times10^{-5}$ | 24.6 |
| | Average | $8.85\times10^{5}$ | $3.67\times10^{-5}$ | 71.3 |
| | 1 | $4.96\times10^{5}$ | $4.83\times10^{-5}$ | 97.5 |
| C-Med | 2 | $1.06\times10^{6}$ | $4.14\times10^{-5}$ | 39.1 |
| | Average | $7.78\times10^{5}$ | $4.49\times10^{-5}$ | 68.3 |

**Table S2:** Kinetic parameters for ImmTAC binding to CD3ε ligand

| **ImmTAC** | **Replicate** | **$k_{on}$ ($M^{-1}s^{-1}$)** | **$K_{off}$ ($s^{-1}$)** | **$K_D$ (nM)** |
|---|---|---|---|---|
| | 1 | $6.18\times10^{4}$ | $1.21\times10^{-2}$ | 196 |
| N-Med | 2 | $5.91\times10^{4}$ | $1.17\times10^{-2}$ | 197 |
| | Average | $6.05\times10^{4}$ | $1.19\times10^{-2}$ | 197 |
| | 1 | $9.09\times10^{3}$ | $9.45\times10^{-2}$ | 9450 |
| N-Lo | 2 | $1.48\times10^{4}$ | $9.30\times10^{-2}$ | 6290 |
| | Average | $1.19\times10^{4}$ | $9.38\times10^{-2}$ | 7870 |
| | 1 | $1.11\times10^{5}$ | $2.19\times10^{-4}$ | 1.97 |
| N-Hi | 2 | $1.36\times10^{5}$ | $2.22\times10^{-4}$ | 1.64 |
| | Average | $1.24\times10^{5}$ | $2.21\times10^{-4}$ | 1.81 |
| | 1 | $4.13\times10^{4}$ | $1.68\times10^{-2}$ | 408 |
| C-Med | 2 | $4.42\times10^{4}$ | $1.72\times10^{-2}$ | 389 |
| | Average | $4.28\times10^{4}$ | $1.70\times10^{-2}$ | 399 |

**Table S3:** Antibody panel used in the mass cytometry experiments

| Extracellular antigen | Metal | Clone | Supplier | RRID |
|---|---|---|---|---|
| cyCD3 | $^{112}Cd$ | UCHT1 | Biolegend | AB_314055 |
| CD4 | $^{143}Nd$ | SK3 | Biolegend | AB_1937277 |
| CD8 | $^{89}Y$ | SK1 | Biolegend | AB_1877104 |
| CD69 | $^{115}In$ | FN50 | Biolegend | AB_2562827 |
| **Intracellular antigen** | | | | |
| Vimentin | $^{111}Cd$ | RV202 | BD Biosciences | AB_393716 |
| CK18 | $^{114}Cd$ | C-04 | SantaCruz | - |
| pPDK1 [S241] | $^{141}Pr$ | J66-653.44.22 | BD Bioscience | AB_647291 |
| cCaspase 3 [D175] | $^{142}Nd$ | D3E9 | CST | AB_10897512 |
| pSTAT5 [Y694] | $^{144}Nd$ | 47/Stat5 | BD Biosciences | AB_2737931 |
| pZAP70/Syk | $^{145}Nd$ | 1503310 | Biolegend | AB_2572026 |
| pSLP76 [Y128] | $^{146}Nd$ | J141-668.36.58 | BD Bioscience | AB_647331 |
| pBTK [Y511] | $^{147}Sm$ | 24a/BTK | BD Bioscience | AB_2067823 |
| pSRC [Y418] | $^{148}Nd$ | SC1T2M3 | BD Bioscience | AB_2865322 |
| p4E-BP1 [T37/46] | $^{149}Sm$ | 236B4 | CST | - |
| pRB [S807/811] | $^{150}Nd$ | J112-906 | BD Biosciences | AB_647295 |
| pNDRG1 [T346] | $^{151}Eu$ | D98G11 | CST | AB_10693451 |
| pAKT [T308] | $^{152}Sm$ | J1-233.371 | BD Biosciences | AB_647259 |
| pCREB [S133] | $^{153}Eu$ | 87G3 | CST | - |
| pSMAD1 [S463/465] | $^{154}Sm$ | D5B10 | CST | - |
| pAKT [S473] | $^{155}Gd$ | D9E | CST | - |
| pNF-kB p65 [S529] | $^{156}Gd$ | K10-895.12.50 | BD Biosciences | AB_647284 |
| pMKK3/MKK6 | $^{157}Gd$ | D8E9 | CST | - |
| pP38 [T180/Y182] | $^{158}Gd$ | D3F9 | CST | - |
| pMAPKAPK2 [T334] | $^{159}Tb$ | 27B7 | CST | - |
| pAMPK$\alpha$ [T172] | $^{160}Gd$ | 40H9 | CST | AB_331250 |
| pBAD [S122] | $^{161}Dy$ | 40A9 | CST | - |
| pHistone H2A.X [S139] | $^{162}Dy$ | D7T2V | CST | AB_2799949 |
| pP90RSK [T359] | $^{163}Dy$ | D1E9 | CST | |
| p120-Catenin [T310] | $^{164}Dy$ | 22/p120 | BD Biosciences | AB_397057 |
| Perforin | $^{165}Ho$ | DG9 | Biolegend | AB_314700 |
| pGSK-3$\beta$ | $^{166}Er$ | D85E12 | CST | - |
| pERK1/2 [T202/Y204] | $^{167}Er$ | 20A | BD Biosciences | AB_399648 |
| pSMAD2 [S465/467] | $^{168}Er$ | D27F4 | CST | - |
| pSTAT3 [Y705] | $^{169}Tm$ | 4/P-Stat3 | BD Biosciences | AB_399646 |
| Granzyme B | $^{170}Er$ | QA16A02 | Biolegend | AB_2686929 |
| pDNAPK [S2056] | $^{171}Yb$ | EPR5670 | Abcam | - |
| pLCK [Y394] | $^{172}Yb$ | A18002D | Biolegend | AB_2814640 |
| pS6 [S240/244] | $^{173}Yb$ | D68F8 | CST | - |
| cPARP [D214] | $^{174}Yb$ | D64E10 | CST | AB_10699459 |
| pHistone H3 [S28] | $^{175}Lu$ | HTA28 | Fluidigm | - |
| Cyclin B1 | $^{176}Yb$ | GNS-11 | BD Biosciences | AB_395290 |
| acHistone H3 [K27] | $^{209}Bi$ | D5E4 | CST | - |

**Table S4:** Summary of the simulation and analysis parameters used in Smoldyn

| | Model 1 | Model 2 | Reference and comment |
|---|---|---|---|
| Close contact radius [μm] | 0.22 | 0.22 | [1] |
| Timestep [s] | 0.002 | 0.002 | [17] |
| Simulation time [s] | 30 | 30 | Larger than scan time |
| TCR diffusion constant [$\mu m^2/s$] | 0.052 | 0.052 | [54] |
| TCR density [$\mu m^{-2}$] | 100 | 100 | [17] |
| pMHC density [$\mu m^{-2}$] | 0, 10 | 0, 10 | No antigen and low antigen |
| ImmTAC binding $k_{on}$ [$\mu m^2/s$] | 0.00748 | 0.00748 | This study |
| ImmTAC unbinding $k_{off}$ [$s^{-1}$] | 0.0119 | 0.0119 | This study |
| pMHC-bound TCR diffusion constant [$\mu m^2/s$] | 0 | 0 | [17] |
| Phosphorylation rate | 0-5 | - | [54] |
| CD45 diffusion constant [$\mu m^2/s$] | - | 0.113 | [54] |
| CD45 density [$\mu m^{-2}$] | - | 600 | [72] |
| CD45 rate constant $k_{CD45}$ [$\mu m^2/s$] | - | 0.001-0.05 | Lower end same order of magnitude as $k_{Lck}$, upper end simulation limit (binding radii cover a significant fraction of simulation area) |
| CD45 exclusion [%] | - | 0-100 | - |
| Lck diffusion constant [$\mu m^2/s$] | - | 0.128 | [54] |
| Lck density [$\mu m^{-2}$] | - | 200 | [72] |
| Lck rate constant $k_{Lck}$ [$\mu m^2/s$] | - | 0.001-0.05 | [54] $k_{cat} = 3.41\ s^{-1}$, at $[Lck] \approx 341\ \mu m^2 \rightarrow k \approx 0.1\ \mu m^2/s$ |
| Scan time [s] | varied around 7.5 | varied around 7.5 | [1] |
